# Nemo-like kinase regulates Progranulin levels in the brain through the microglial endocytosis-lysosomal pathway

**DOI:** 10.1101/358010

**Authors:** Tingting Dong, Hiroshi Kokubu, Terri M. Driessen, Leon Tejwani, Janghoo Lim

**Affiliations:** Department of Genetics, Yale School of Medicine, New Haven, USA; Interdepartmental Neuroscience Program, Yale University, New Haven, USA; Department of Neuroscience, Yale School of Medicine, New Haven, USA; Program in Cellular Neuroscience, Neurodegeneration and Repair, Yale School of Medicine, New Haven, USA

## Abstract

Genetic variants in *Granulin (GRN)*, which encodes the secreted glycoprotein Progranulin (PGRN), are associated with several neurodegenerative diseases including frontotemporal lobar degeneration, neuronal ceroid lipofuscinosis, and Alzheimer’s disease. These genetic alterations manifest in pathological changes due to a reduction of PGRN expression; therefore, identifying a factor that can modulate PGRN levels *in vivo* would enhance our understanding of PGRN in neurodegeneration, and could reveal novel potential therapeutic targets. Here, we report that Nemo-like kinase (Nlk) regulates Pgrn levels and its associated neuropathophysiology. Genetic interaction studies in mice show that *Grn* heterozygote mice on an *Nlk* heterozygote background display pathological and behavioral phenotypes which mimic *Grn* knockout mice. Furthermore, biochemical and cell biological studies suggest that Nlk reduction promotes Pgrn degradation via the endocytosis-lysosomal pathway, specifically in microglia. Our results reveal a new mechanism for the regulation of Pgrn in the brain and provide insight into the pathophysiology of PGRN-associated diseases.

## Introduction

Progranulin (PGRN) is an evolutionarily conserved, cysteine-rich, secreted glycoprotein encoded by the Granulin (*GRN*) gene (Cenik et al., 2012; Toh et al., 2011). In humans, several neurodegenerative diseases are closely linked to the reduction of PGRN expression. Haploinsufficiency of PGRN, caused by heterozygous loss-of-function mutations in one *GRN* allele, leads to the development of familial frontotemporal lobar degeneration (FTLD), a neurodegenerative disease characterized by atrophy in the frontal and temporal lobes of the brain (Baker et al., 2006; Cenik et al., 2012; Chen-Plotkin et al., 2010; Cruts et al., 2006; Gijselinck et al., 2008). Homozygous loss-of-function mutations in both alleles of *GRN* cause neuronal ceroid lipofuscinosis (NCL), a neurodegenerative disease in which lipofuscins aberrantly accumulate in the affected tissues (Smith et al., 2012). Finally, specific single nucleotide variants in *GRN* have been shown to decrease plasma and brain expression levels of PGRN, and are risk factors for Alzheimer’s disease (AD), the most common form of dementia (Perry et al., 2013; Takahashi et al., 2017; Wojtas et al., 2012).

Although the involvement of *GRN* genetic variants in human disease appears to be straightforward, the precise effect of changes to *Grn* in animal models is far more elusive. In mice, *Grn* heterozygosity causes little or no change in behavior and neuropathology up to 23 months of age (Ahmed et al., 2010). In contrast, mice lacking both copies of *Grn* display many behavioral and neuropathological abnormalities, which recapitulate several features of PGRN-deficient FTLD (FTLD-PGRN) and NCL (Filiano et al., 2013; Hafler et al., 2014; Klein et al., 2017; Yin et al., 2010b). In addition, Pgrn has been shown to be a modulator in animal models of neurodegenerative diseases including AD (Minami et al., 2014; Takahashi et al., 2017) and Parkinson’s disease (Van Kampen et al., 2014). Together these studies suggest that decreased expression of PGRN is a direct cause, or indirect risk factor, for a variety of neurodegenerative diseases.

The mechanism through which reductions in PGRN expression lead to diverse neurodegeneration phenotypes remains a largely unanswered question in the field. Pgrn is a secreted glycoprotein which is endocytosed through receptor- and clathrin-dependent endocytosis and delivered to the endosome/lysosome pathway. Previous works suggest that Pgrn plays important roles in the regulation of neurite growth, innate immunity, and lysosome function, all of which are cellular processes studied extensively in the context of neurodegeneration (Ahmed et al., 2007; Gass et al., 2012; Kessenbrock et al., 2008; Klein et al., 2017; Lui et al., 2016; Tanaka et al., 2017; Van Damme et al., 2008). Loss of Pgrn in neurons results in the inhibition of neurite outgrowth (Gass et al., 2012; Van Damme et al., 2008), and Pgrn is involved in the modulation of Wnt and Notch signaling pathways, which regulate axon regeneration, neuronal differentiation, and neuronal survival (Altmann et al., 2016; Raitano et al., 2015; Rosen et al., 2011). In addition, it has been shown that Pgrn plays a role in the regulation of neuroinflammation as an anti-inflammatory factor (Ahmed et al., 2007; Kessenbrock et al., 2008). Lysosomal defects and excessive complement production, which triggers selective synaptic pruning by microglia, occur with loss of Pgrn (Lui et al., 2016). Several other studies also suggest that Pgrn plays active roles in the control of lysosome formation and function (Klein et al., 2017; Tanaka et al., 2017).

Besides its role in the regulation of neurite outgrowth, neuroinflammation, and lysosomes, several studies have focused on Pgrn catabolism. Pgrn is expressed in a wide range of tissues and cell types, including neurons and activated microglia in the brain (Hu et al., 2010; Petkau et al., 2010). It undergoes receptor-mediated endocytosis and subsequent trafficking to the lysosomes. Extracellular Pgrn can be endocytosed through a Sortilin receptor-dependent or - independent manner and degraded by endosome-mediated delivery to lysosomes (Hu et al., 2010; Zhou et al., 2015). Thus, identifying factors capable of modulating the functioning and/or expression levels of Pgrn itself will be critical, as knowledge of such regulatory factors will not only impart invaluable information regarding the underlying pathophysiology of PGRN-associated neurodegenerative diseases, but also uncover novel therapeutic targets.

Nemo-like kinase (Nlk) is an evolutionarily conserved serine/threonine kinase that plays roles in various signaling pathways including Notch, Wnt, and DNA damage response pathways (Ishitani et al., 1999; Meneghini et al., 1999; Ota et al., 2012; Rocheleau et al., 1999; Rottinger et al., 2006; Zhang et al., 2014a). Several lines of evidence suggest that Nlk could act as a potential key protein to modulate the pathogenesis of PGRN-associated neurological diseases and the expression levels of PGRN in the brain. First, changes in Nlk and Pgrn levels affect similar cellular phenotypes. For example, both Nlk and Pgrn are known to be important for neurite outgrowth (Gass et al., 2012; Ishitani et al., 2009; Van Damme et al., 2008) and for the regulation of neuroinflammation (Ahmed et al., 2007; Cenik et al., 2012; Kessenbrock et al., 2008; Saijo et al., 2009). Second, Nlk and Pgrn influence similar molecular pathways that may underlie shared cellular functions. Both Nlk and Pgrn play a role in the modulation of Wnt and Notch signaling pathways, which are important for neuronal survival and axon regeneration (Altmann et al., 2016; Ishitani et al., 1999; Meneghini et al., 1999; Ota et al., 2012; Raitano et al., 2015; Rocheleau et al., 1999; Rosen et al., 2011; Rottinger et al., 2006). Third, and most importantly, deletion of a component in the Nlk-mediated molecular signal transduction pathways leads to neuropathological phenotypes that are similar to AD and FTLD-PGRN (Christopher et al., 2017). Specifically, Nlk-mediated phosphorylation of Nurr1 is essential for the prevention of neuroinflammation by inhibiting neurotoxic gene expression in microglia via the recruitment of CoREST/LSD1 complex for the transrepression of NF-kB activity (Saijo et al., 2009). Subsequently, mice lacking the histone demethylase *Lsd1* display behavioral, neuropathological, and molecular phenotypes that highly overlap with those seen in human AD and FTLD-PGRN cases (Christopher et al., 2017).

In this study, we investigated the potential role of Nlk in modulating Pgrn expression and function by using mouse models and cultured cells. Our genetic interaction studies in mice demonstrate that *Grn* heterozygote mice display neuropathological and behavioral abnormalities in an *Nlk* heterozygote background at 1 year of age, whose phenotypes are similar to those previously observed in *Grn* knockout mice. The phenotypes observed in the compound heterozygote arise due to Nlk’s function as a positive regulator of brain Pgrn levels, as we show that loss of Nlk leads to a reduction in Pgrn levels. Furthermore, Nlk-mediated regulation of Pgrn levels in the brain occurs mainly through microglia by controlling the clathrin-dependent Pgrn endocytosis and degradation via lysosomes. These results reveal a new mechanism for the regulation of Pgrn levels in the brain and provide insight into the pathophysiology of several neurodegenerative diseases associated with PGRN expression levels, such as AD, FTLD, and NCL.

## Results

### Decreased *Nlk* expression induces neuropathological phenotypes in heterozygous *Grn* mice

Previous studies have documented that *Grn^+/−^* heterozygous mice do not display any obvious pathological and behavioral phenotypes up to 23 months of age (Ahmed et al., 2010). In contrast, *Grn^−/−^* null mice exhibit mild to moderate cognitive impairment and pathological abnormalities at 12-18 months of age (Ghoshal et al., 2012; Petkau and Leavitt, 2014; Yin et al., 2010b). Although *Grn^+/−^* mice are phenotypically normal, it is possible that *Grn^+/−^* mice may exhibit a synergistic phenotype in certain genetic backgrounds, such as with reduced Nlk expression. To explore the possibility that the neuropathological phenotypes associated with loss of *Grn* are affected by Nlk, we conducted a genetic interaction study using mouse models of *Grn* and *Nlk*. Since both genes are located on the same chromosome, we crossed *Grn^+/−^* mice with *Nlk^+/−^*animals and evaluated the progeny on a pure C57BL/6J background, investigating whether *Nlk^+/−^Grn^+/−^* double heterozygote mice could develop pathological and behavioral changes.

We first investigated whether a 50% reduction of Nlk expression generates pathological phenotypes in *Grn^+/−^* mice. Previous studies have demonstrated that *Grn^−/−^* mice recapitulate several key features of PGRN-FTLD and NCL, including microgliosis, accumulation of lipofuscin, and retinal degeneration (Ahmed et al., 2010; Filiano et al., 2013; Hafler et al., 2014; Klein et al., 2017; Yin et al., 2010a). We therefore examined these phenotypes in 1-year-old *Nlk^+/−^ Grn^+/−^* mice to determine if loss of *Nlk* promotes pathology in the phenotypically normal *Grn^+/−^* mice.

To identify changes in microgliosis, Iba1 and CD68 antibodies labeling microglia and microglial lysosomes, respectively, were used for immunohistochemistry. There was a significant increase in the number of Iba1-positive cells in the thalamus of 1-year-old *Nlk^+/−^ Grn^+/−^* mice compared to age-matched littermate controls (Figure 1A,B). In addition, CD68-positive immunostaining showed enlarged lysosome volume in Iba1-positive microglia in 1-year-old *Nlk^+/−^ Grn^+/−^* mice (Figure 1A,C). For both Iba-1 and CD68-positive immunostaining, there was no significant difference between *Nlk^+/−^* and *Grn^+/−^* mice with wild-type (WT) control littermates (Figure 1A-C), further indicating that partial loss of *Nlk* and *Grn* contributes to microgliosis. These results are consistent with a previous report that showed microglia activation in *Grn^−/−^* mice (Lui et al., 2016). To examine lipofuscin deposition, autofluorescence was assessed in the retina, thalamus, and cortex in 1-year-old mice. There was a significant accumulation of lipofuscin in the retina of *Nlk^+/−^ Grn^+/−^* mice compared to littermate controls (Figure 1D,E), and a trend towards lipofuscin accumulation in the thalamus (Figure 1-figure supplement 1A-C) (p=0.1522 *one-way ANOVA* with Tukey’s multiple comparisons post-hoc test) and cortex (Figure 1-figure supplement 1D,E) (p=0.1381 *one-way ANOVA* with Tukey’s multiple comparisons post-hoc test), which were significant with the non-parametric Mann-Whitney *t-test* (P<0.05). To explore retinal degeneration in this mouse model, immunohistochemistry was conducted for Brn3a, a transcription factor expressed in most retinal ganglion cells. We found substantial degeneration of retinal ganglion cells in 1-year-old *Nlk^+/−^ Grn^+/−^* mice (Figure 1F,G). Taken together, these neuropathological results show that Nlk may play an important role in modulating phenotypes typified by Pgrn reduction.

**Figure 1.**
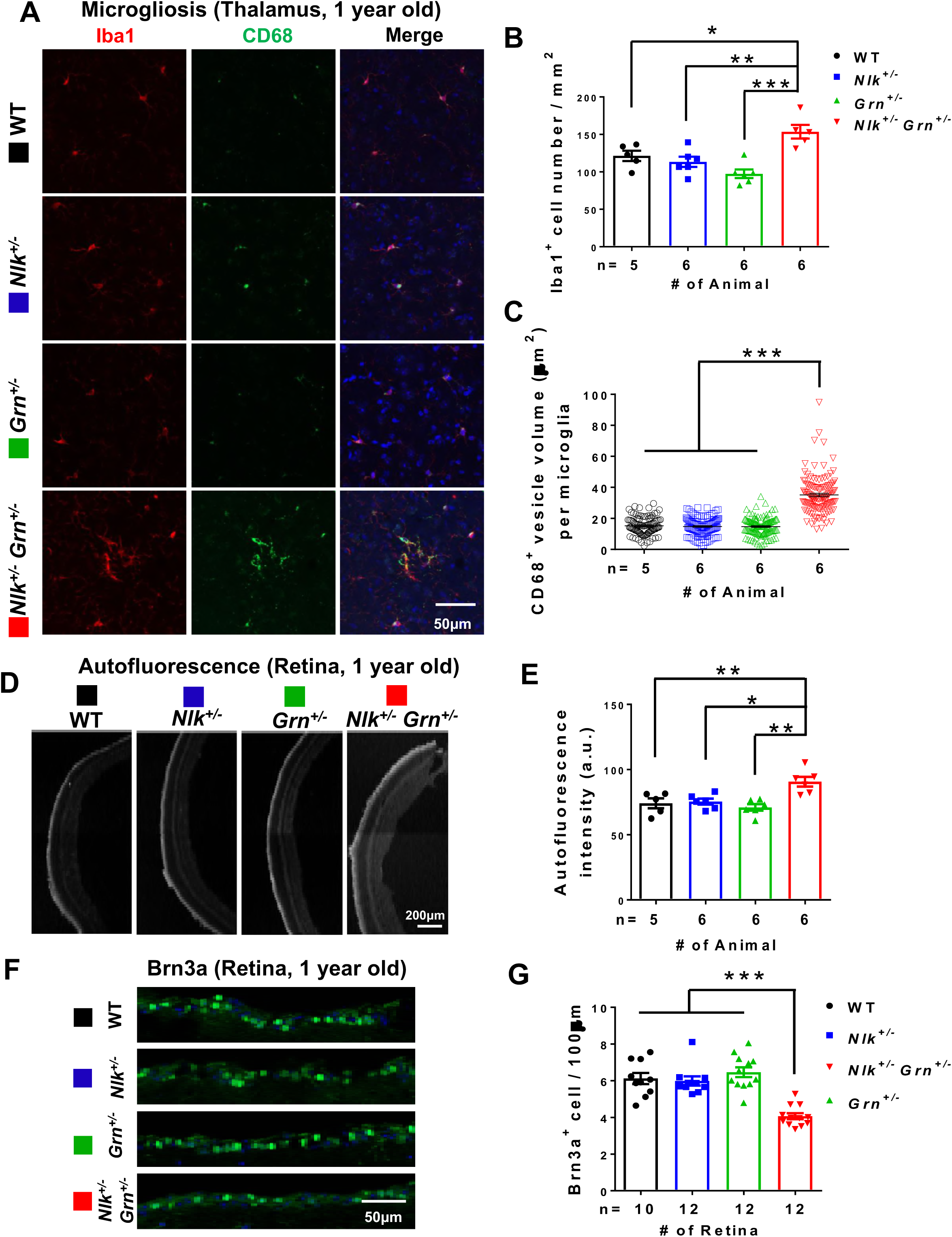
*Nlk^+/−^ Grn^+/−^* mice display pathological phenotypes that are similar to those seen in *Grn^−/−^*. **(A-C)** Increased microglial activation in *Nlk^+/−^ Grn^+/−^* mice compared to their littermates. Representative confocal images **(A)** and quantification **(B,C)** of Iba1 and CD68 staining from the thalamus of 1-year-old mice. **(B)** The number of Iba1-positive microglia was dramatically increased in the thalamus of *Nlk^+/−^ Grn^+/−^* mice. Error bars represent standard error of the mean (SEM) in this and all following graphs. *P < 0.05, **P < 0.01, ***P < 0.001 (*one-way ANOVA* with Tukey’s multiple comparisons post-hoc test). **(C)** CD68-positive vesicle volume in Iba1-positive microglia was also significantly increased in *Nlk^+/−^ Grn^+/−^* mice. ***P < 0.001 (*one-way ANOVA* with Tukey’s multiple comparisons post-hoc test). **(D,E)** Representative images **(D)** and quantification **(E)** of autofluorescence using 488 nm excitation in the retina of 1-year-old WT, *Nlk^+/−^*, *Grn^+/−^*, *Nlk^+/−^ Grn^+/−^* mice. *p < 0.05, **p < 0.01 (*one-way ANOVA* with Tukey’s multiple comparisons post-hoc test). **(F,G)** Representative images **(F)** and quantification **(G)** of 1-year old WT, *Nlk^+/−^*, *Grn^+/−^*, *Nlk^+/−^Grn^+/−^* mouse retinas stained for Brn3a (Green) and DAPI (blue). ***p < 0.001 (*one-way ANOVA* with Tukey’s multiple comparisons post hoc-test). In this and all following figures, mouse genotypes are color-coded. Black lines or bars represent WT, blue for *Nlk^+/−^*, green for *Grn^+/−^*, and red for *Nlk^+/−^ Grn^+/−^* unless otherwise mentioned.

### *Nlk* reduction promotes behavioral alterations in phenotypically unaffected *Grn^+/−^* mice

We next examined the effect of *Grn^+/−^ Nlk^+/−^* compound heterozygosity on mouse behavior. A recent report has documented that *Grn^−/−^* deficient mice show disinhibition in the context of the elevated plus maze test (Klein et al., 2017), which is in agreement with FTLD patient behavioral changes (de Vugt et al., 2006). To determine if the genetic interaction between *Nlk* and *Grn* induces this behavioral phenotype in the otherwise normal *Grn^+/−^* mice, we assessed 1-year-old *Grn^+/−^ Nlk^+/−^* mice performance on the elevated plus maze (Figure 2A). Since previous reports have found genotype-dependent alterations in levels of anxiety specifically in male *Grn^−/−^* mice, only males were tested (Kayasuga et al., 2007; Petkau et al., 2012). *Nlk^+/−^ Grn^+/−^* male mice were less anxious and disinhibited, as they spent more time in the open arms compared to littermate controls (Figure 2A). *Grn^+/−^* and *Nlk^+/−^* mice alone did not have any significant behavioral phenotype relative to WT controls (Figure 2A).

**Figure 2.**
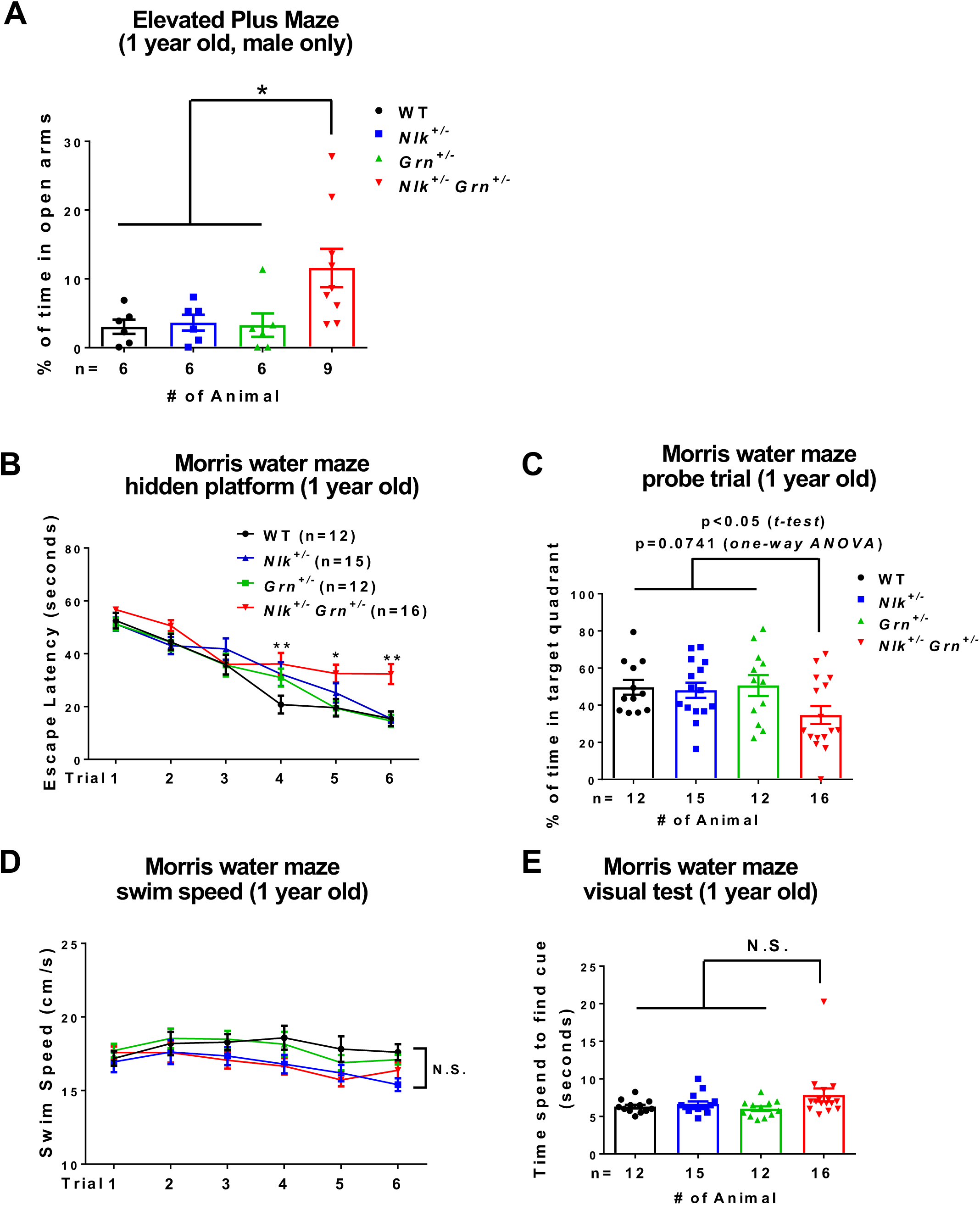
*Nlk^+/−^ Grn^+/−^* double heterozygote mice display *Grn^−/−^-like* behavioral deficits. **(A)** *Nlk^+/−^ Grn^+/−^* male mice showed decreased anxiety in elevated plus maze test by spending significantly longer time in the open arms. *P < 0.05 (*one-way ANOVA* with Tukey’s multiple comparisons post-hoc test). **(B-E)** In the Morris water maze (MWM) test, **(B)** *Nlk^+/−^ Grn^+/−^* mice showed significantly prolonged escape latency in the hidden-platform test. *P < 0.05, **P < 0.01 (repeated measures *two-way ANOVA* with Tukey’s HSD test relative to WT). **(C)** *Nlk^+/−^ Grn^+/−^* mice also spent shorter time in the trained target quadrant during probe trial one day after the three-day training period. P=0.0741 (*one-way ANOVA* with Tukey’s multiple comparisons post-hoc test); *P < 0.05 (non-parametric Mann-Whitney *t-test).* **(D,E)** There is no significant difference in swimming speed in the hidden-platform test **(D)** or in visual ability by visual platform test **(E).** N.S., non-significant (repeated measures *two-way ANOVA* with Tukey’s HSD test for **D,** *one-way ANOVA* with Tukey’s multiple comparisons post-hoc test for **E).**

We also examined cognitive behavior by performing a Morris water maze (MWM) test at 1 year of age. *Nlk^+/−^ Grn^+/−^* mice had a significant impairment in identifying the hidden platform on trials 4-6 of testing relative to WT controls (Figure 2B). As expected, *Grn^+/−^* mice performed similarly to WT controls (Figure 2B). During the probe trial, *Grn^+/−^ Nlk^+/−^* mice spent less time in the target quadrant than their WT littermates (Figure 2C) (p=0.0741, *one-way ANOVA* with Tukey’s multiple comparisons post-hoc test), which was significant with non-parametric Mann Whitney *t-test* (P< 0.05). To verify that these results were not an effect of alterations in locomotor behavior and visual acuity, swim speed, and vision were also assessed (Figure 2D,E). There was no significant difference between *Nlk^+/−^ Grn^+/−^* mice and their littermates in swim speed across all six trials of MWM testing (Figure 2D). Although we observed significant retinal degeneration in *Nlk^+/−^ Grn^+/−^* mice (Figure 1F,G), this did not produce an observable behavioral deficit in vision, as there was no difference in the time spent to identify a visual cue (Figure 2E). Collectively, these data indicate that *Nlk^+/−^ Grn^+/−^* mice have normal sensorimotor function. These results are correlated to the observed trending pathological phenotypes (Figure 1), as the thalamus, at least in part, contributes to the anxiety phenotypes and the cortex is important for the recognition phenotype. In addition, consistent with our pathological studies, these behavioral assays support the hypothesis that partial loss of *Nlk* and *Grn* additively or synergistically contributes to the appearance of pathological and behavioral phenotypes found in *Grn^−/−^* mice.

### Decreased expression of *Nlk* reduces Pgrn levels *in vivo*

We next investigated the molecular basis by which *Nlk* reduction in *Grn^+/−^* mice promotes the appearance of pathological and behavioral phenotypes. Since *Nlk^+/−^ Grn^+/−^* mice show phenotypes similar to *Grn^−/−^* mice, we wondered whether partial loss of *Nlk* decreases the expression levels of Pgrn. We therefore measured both intracellular and extracellular Pgrn expression, since it is a secreted protein. We found that the expression levels of both intracellular and extracellular Pgrn proteins were significantly reduced in the cortex of 1-year-old *Nlk^+/ −^Grn^+/−^* mice compared to *Grn^+/−^* littermates (Figure 3A-C). Interestingly, *Nlk^+/−^* mice alone had a significant reduction in extracellular Pgrn expression relative to WT control littermates, with comparable Pgrn expression between *Nlk^+/−^* and *Grn^+/−^* mice (Figure 3A-C). Immunofluorescence staining for Pgrn in the mouse cortex confirmed our western blotting results, showing a significant decrease in Pgrn expression in *Nlk^+/−^ Grn^+/−^* relative to *Grn^+/−^,* and a decrease in *Nlk^+/−^*relative to WT controls (Figure 3D,E). These data suggest that Nlk positively regulates Pgrn expression *in vivo.*

**Figure 3.**
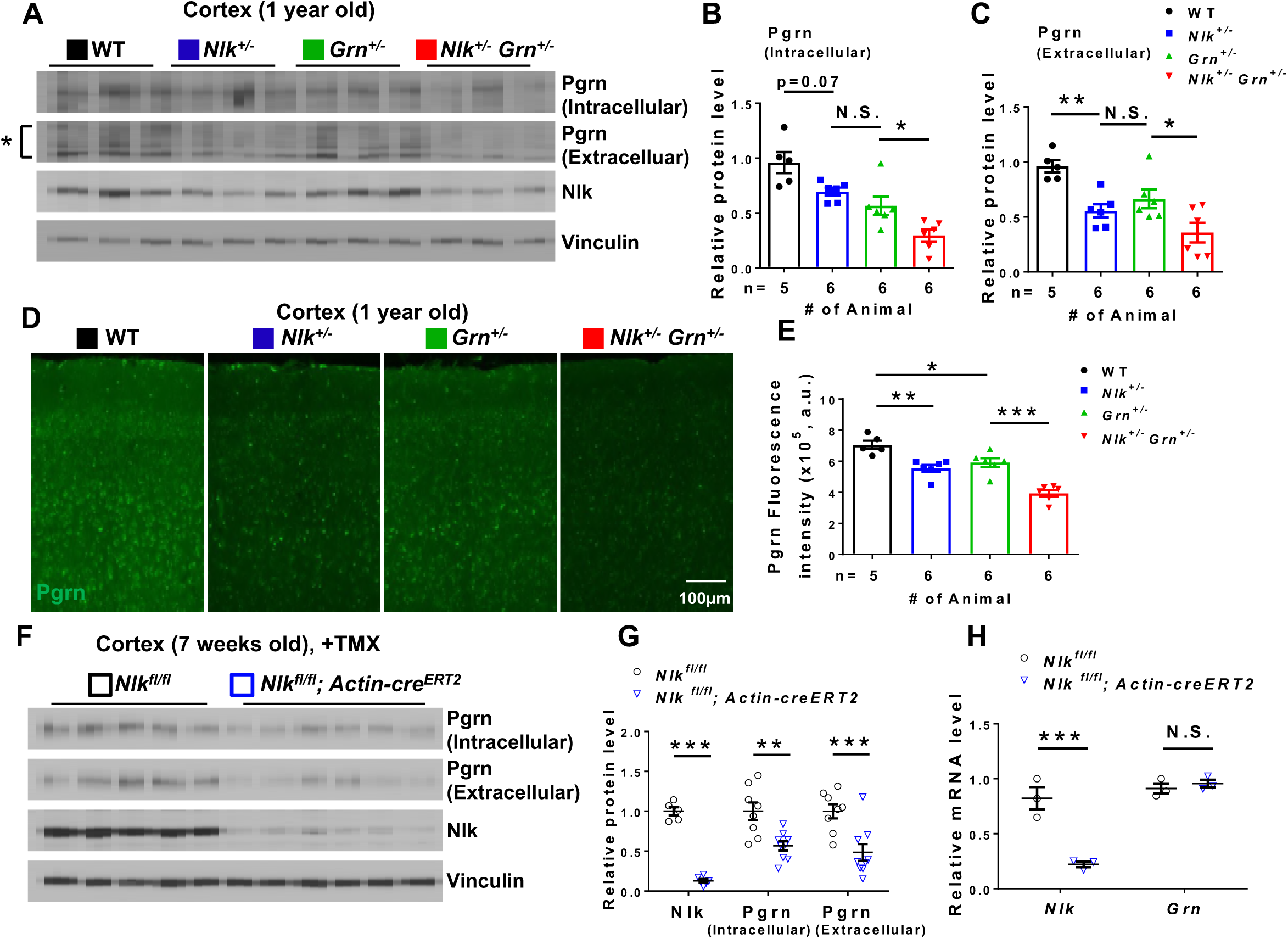
Loss of Nlk regulates Pgrn levels in the mouse cortex. **(A-E)** Expression levels of Pgrn were significantly decreased in the mouse cortex of *Nlk^+/−^ Grn^+/−^* mice compared to their littermate controls. Representative western blot images **(A)** and quantification **(B,C)** of Pgrn and Nlk expression in the 1-year-old mouse cortex. Normalized protein levels of Nlk and Pgrn signals to Vinculin are shown in this and all following graphs. *P < 0.05, **P < 0.01, N.S., non-significant (*one-way ANOVA* with Tukey’s multiple comparisons post-hoc test). Representative confocal images **(D)** and quantification **(E)** of Pgrn immunofluorescent staining in a 1-year-old mouse cortex. Total fluorescence intensity quantification across the image field was automated using Volocity software. *P < 0.05, **P < 0.01, ***P < 0.001 (*one-way ANOVA* with Tukey’s multiple comparisons post-hoc test). **(F-H)** Temporal deletion of Nlk in all cell types reduces Pgrn protein levels in the mouse cortex. Protein and mRNA expression levels were analyzed in the 7-weeks-old cortex of *Nlk^fl/fl^* and *Nlk^fl/fl^;Actin-creERT2* mice after tamoxifen (TMX) injection. Representative western blot images **(F)** and quantification **(G)** showing the reduced Pgrn expression and Nlk in the mouse cortex. **P < 0.01, and ***P < 0.001 (non-parametric Mann-Whitney *t-test,* n=8 animals for *Nlk^fl/fl^*, n=9 for *Nlk^fl/fl^; Actin-creERT2).* **(H)** Quantification of *Grn* mRNA expression levels in *Nlk* deleted mouse cortex, showing no transcriptional effects. Normalized levels of *Nlk* and *Grn* mRNA to mouse *ACTB* are shown in this and all following graphs. ***P < 0.001, N.S., non-significant (non-parametric Mann-Whitney *t-test,* n=3 per group).

To determine if a complete loss of *Nlk* enhances the down-regulation of Pgrn, we temporally deleted *Nlk* in all cell types during adulthood to circumvent *Nlk* knockout (KO) prenatal lethality (Ju et al., 2013). We took advantage of mice with a flox-*Nlk* allele (Canalis et al., 2014) and crossed them to mice expressing tamoxifen (TMX)-inducible Cre driven ubiquitously by a CMV-IE enhancer and chicken ß-actin promoter (Hayashi and McMahon, 2002). We generated *Nlk^fl/fl^*; *Actin-cre^ERT2^* mice and their littermate mice on a pure C57BL/6J background. After intraperitoneal TMX injection for 7 consecutive days starting at 6 weeks old, we verified that the expression levels of *Nlk* mRNA and protein were significantly reduced in the cortex of *Nlk^fl/fl^*; *Actin-cre^ERT2^* mice (Figure 3F-H). Furthermore, there was a significant reduction of intracellular and extracellular Pgrn protein levels in the cortex of *Nlk^fl/fl^*; *Actin-cre^ERT2^* mice compared to their littermates (Figure 3F,G).

The down-regulation of Pgrn protein expression in *Nlk^fl/fl^*; *Actin-cre^ERT2^* mice may be due to a similar down-regulation at the transcriptional level. To determine if *Grn* mRNA was also down-regulated in *Nlk^fl/fl^;Actin-cre^ERT2^* mice, we utilized quantitative real-time reverse transcription polymerase chain reaction (RT-qPCR) to assess *Grn* expression in the cortex of 7-week-old mice. *Grn* mRNA level was not significantly altered by *Nlk* deletion (Figure 3H). Together, these studies indicate that Nlk regulates Pgrn expression in the adult brain post-transcriptionally.

### Nlk regulates Pgrn levels via microglia, but not through neurons

Since we have observed that constitutive deletion of a single *Nlk* allele (Figure 3A-E) or temporally induced deletion of both copies of *Nlk* in all cell types (Figure 3F-H) leads to a significant reduction of Pgrn protein expression in the adult mouse cortex, we next wanted to determine which cell types are essential for the Nlk-mediated regulation of Pgrn expression. *Nlk* and *Grn* are expressed in both neurons and glia in the cortex (Zhang et al., 2014b), so we first tested whether Nlk-mediated regulation of Pgrn primarily occurs in neurons, and generated *Nlk^fl/fl^*; *Nex-cre* mice and their littermate controls on a pure C57BL/6J background. *Nex-cre* mice express Cre recombinase in the principal neurons of the cortex and the hippocampus and persists through adulthood (Goebbels et al., 2006). The expression levels of *Nlk* mRNA and protein were significantly reduced in the cortex of *Nlk^fl/fl^; Nex-cre* mice at 6 weeks of age (Figure 4A-C). However, we failed to find any significant alterations in the expression levels of Pgrn protein or its mRNA in the cortex of *Nlk^fl/fl^; Nex-cre* mice *in vivo* (Figure 4A-C).

**Figure 4.**
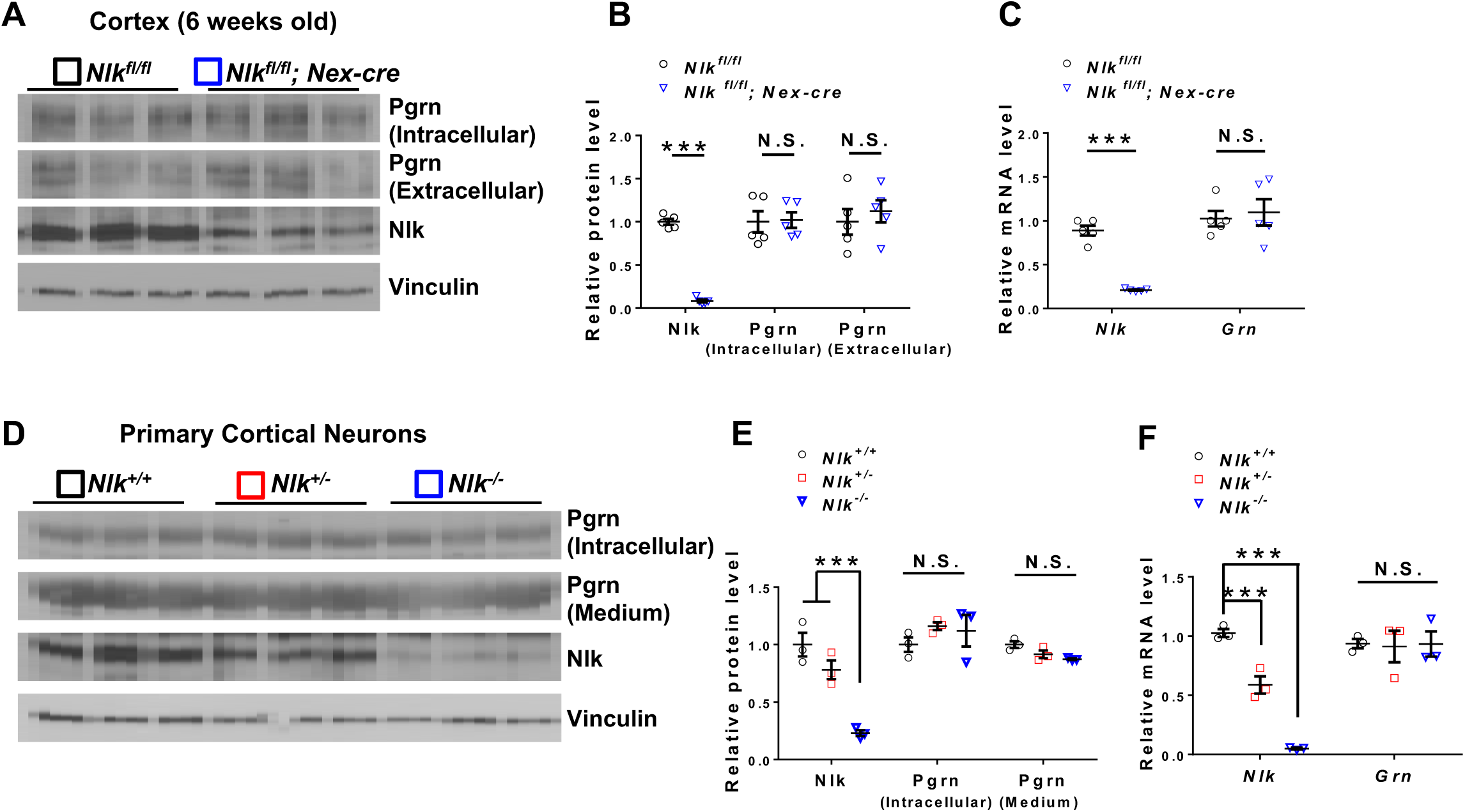
Nlk does not regulate Pgrn levels in neurons. **(A-C)** Nlk does not regulate Pgrn levels through neurons *in vivo.* Protein and mRNA expression levels were analyzed in the 6-weeks-old cortex of *Nlk^fl/fl^* and *Nlk^fl/fl^; Nex-cre* mice. Representative western blot images **(A)** and quantification **(B,C)** showing that the expression levels of Pgrn protein or *Grn* mRNA in the whole cortex were not altered by Nlk deletion specifically in the principal neurons. ***P < 0.001, N.S., non-significant (non-parametric Mann-Whitney *t-test,* n=5 animals per group). **(D-F)** Nlk does not regulate Pgrn levels in primary neurons *in vitro.* Expression levels of Pgrn protein and *Grn* mRNA were analyzed in cultured primary cortical neurons from WT, *Nlk^+/−^,* and *Nlk^−/−^* mice. Representative western blot images **(D)** and quantification **(E,F)** showing that the expression levels of Pgrn (intracellular and extracellular/medium) or *Grn* mRNA were not altered by Nlk expression levels in neurons. ***P < 0.001, N.S., non-significant (*one-way ANOVA* with Tukey’s post-hoc testing, n=3 per group).

Since *Nex-cre* is expressed mainly in excitatory neurons, but not in inhibitory interneurons (Goebbels et al., 2006), we assayed Pgrn expression in constitutive *Nlk^+/−^* and *Nlk^−/−^*primary neuron cortical culture, which contains both excitatory and inhibitory neurons (Lodato and Arlotta, 2015). There was no significant difference in Pgrn protein or mRNA expression levels between WT, *Nlk^+/−^,* and *Nlk^−/−^* cortical neurons *in vitro* (Figure 4D-F). Taken together, these data suggest that the Nlk-mediated Pgrn regulation observed *in vivo* may not occur in neurons.

Transcriptomics studies have indicated that *Grn* is expressed highly in microglia (Zhang et al., 2014b). To test if microglia could be the primary cell type in Nlk-mediated Pgrn regulation, we generated *Nlk^fl/fl^; Cx3cr1-cre* mice and their littermate mice on a pure C57BL/6J background. *Cx3cr1-cre* mice express Cre recombinase in microglia in the brain (Yona et al., 2013). We failed to detect a statistically significant decrease in Nlk expression in the whole cortex of *Nlk^fl/fl^; Cx3cr1-cre* mice (Figure 5A-C), which is likely due to the low fraction of microglia and therefore, minor contribution of Nlk expression from microglia in the whole mouse cortex. Furthermore, we are unable to detect the reduction or loss of Nlk in microglia by immunostaining due to non-specific signals. However, we found that the expression levels of intracellular and extracellular Pgrn protein were significantly reduced in *Nlk^fl/fl^; Cx3cr1-cre* mice compared to littermate controls at 6 weeks of age (Figure 5A,B). Consistent with results from *Nlk^fl/fl^*; *Actin-cre^ERT2^* mice (Figure 3H), there was no significant change in *Grn* mRNA in *Nlk^fl/fl^*; *Cx3cr1-cre* mice compared to littermate controls (Figure 5C). To further confirm the role of microglia in Nlk-mediated regulation of Pgrn levels, we also performed primary microglia culture experiments from the cortex of WT and *Nlk^+/−^* mice and monitored Pgrn expression. Consistent with our finding *in vivo, Nlk* haploinsufficiency resulted in a marked reduction of Pgrn expression in primary microglia culture *in vitro* (Figure 5D,E).

**Figure 5.**
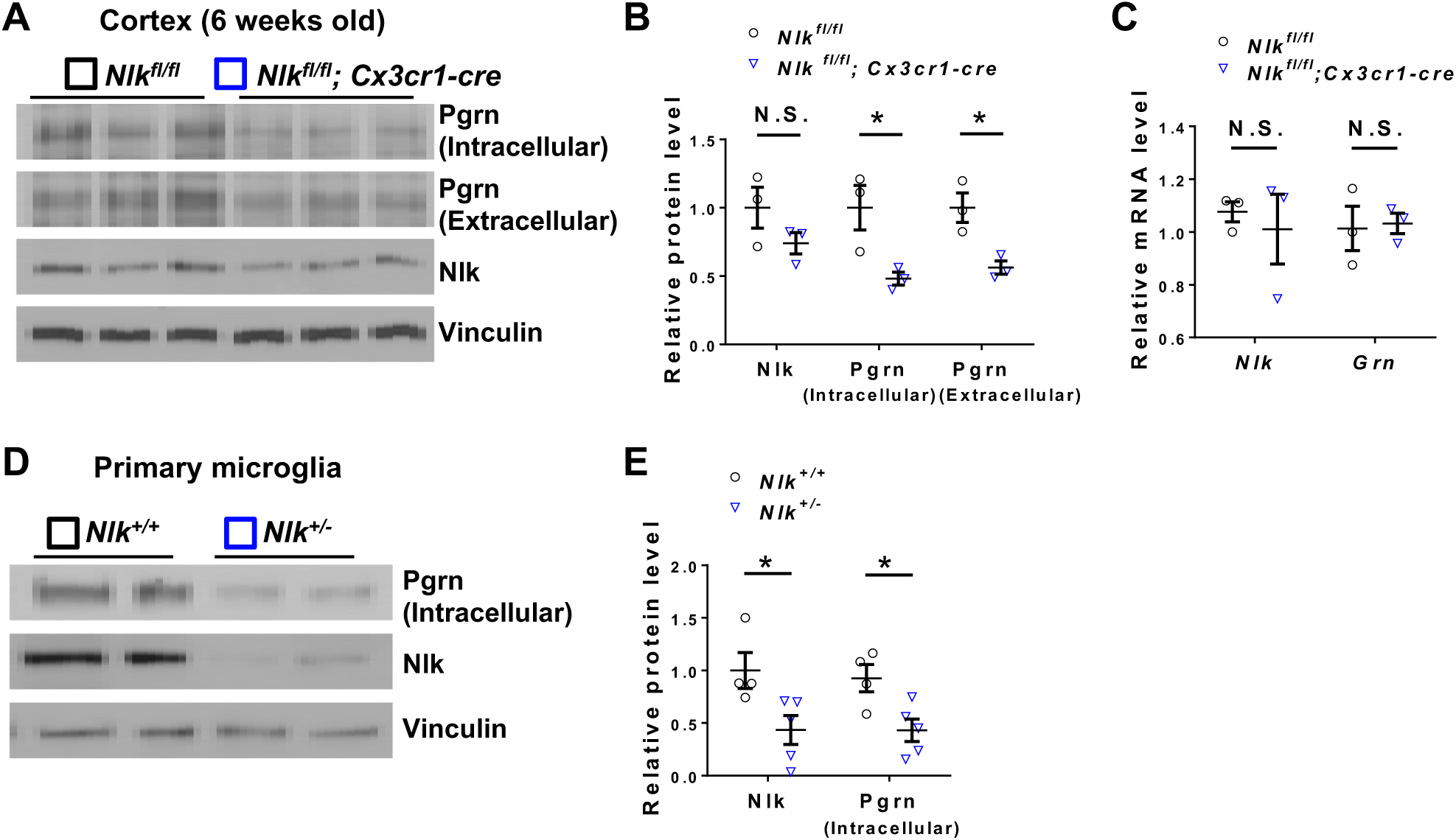
Nlk regulates Pgrn levels in microglia. **(A-C)** Nlk regulates Pgrn levels in microglia *in vivo.* Protein and mRNA expression levels were analyzed in the 6-weeks-old cortex of *Nlk^fl/fl^*and *Nlk^fl/fl^; Cx3cr1-cre* mice. Representative western blot images **(A)** and quantification **(B)** showing that the protein expression levels of Pgrn are significantly decreased by Nlk deletion in microglia. *P < 0.05, N.S., non-significant (non-parametric Mann-Whitney *t-test,* n=3 animals per group). Quantification of *Grn* mRNA expression levels in the mouse cortex from microglia-specific *Nlk* deletion mice **(C).** N.S., non-significant (non-parametric Mann-Whitney *t-test,* n=3 per group). **(D,E)** Reduced Nlk expression significantly decreased Pgrn expression levels in primary microglia. Representative western blot images **(D)** and quantification **(E)** of Pgrn expression levels in primary microglia at DIV-16 from *Nlk^+/+^* and *Nlk^+/−^* mice. *P < 0.05 (non-parametric Mann-Whitney *t-test,* n=5 per group).

Since loss of *Nlk* mediates a down-regulation of Pgrn, increasing *Nlk* expression should in turn elevate Pgrn expression. To test this, we utilized a murine microglial cell line, BV2. To first verify that BV2 cells recapitulate Nlk-mediated regulation of Pgrn as seen in primary microglia, we intended to generate Nlk-KO cells using CRISPR/Cas9 (Cong et al., 2013). Due to the additional chromosomes that result from the non-diploid nature of the BV2 cancer cell line, incomplete *Nlk* gene-targeted BV2 cell clones were identified and one clone was selected as Nlk knockdown (Nlk-KD) cells for our analyses, which led to a consistent knockdown efficiency of Nlk (about 70% reduction) at the protein level (Figure 6A,B) and a significant reduction at the mRNA level (Figure 6C). Consistent with our results from primary microglia (Figure 5D,E), there was a significant reduction in intracellular and medium/extracellular Pgrn in *Nlk*-KD BV2 cells (Figure 6A,B), and no change in *Grn* mRNA (Figure 6C), validating BV2 cells as a model for identifying the mechanism underlying Nlk’s moderating effect on Pgrn levels.

**Figure 6.**
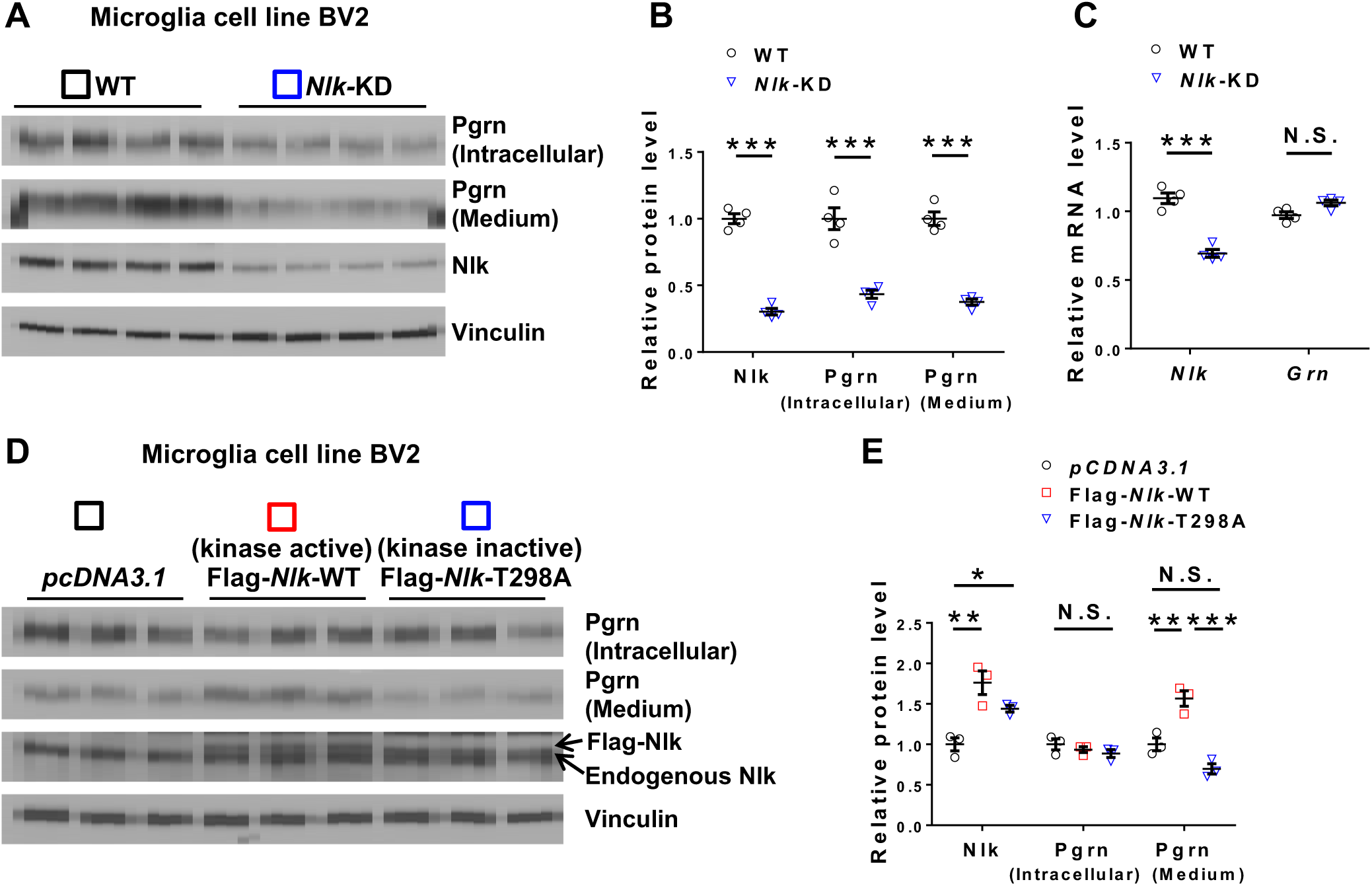
Nlk regulates Pgrn levels in a kinase activity-dependent manner in BV2 microglial cells. **(A-C)** Nlk reduction resulted in decreased Pgrn levels in BV2 microglial cells. Protein and mRNA expression levels were analyzed in cultured microglia cell lines from WT and Nlk-KD BV2 cells. Representative western blot images **(A)** and quantification **(B,C)** showing the expression levels of Pgrn and Nlk proteins and their corresponding mRNAs. ***P < 0.001, N.S., non-significant (non-parametric Mann-Whitney *t-test,* n=4 per group). **(D,E)** Increased expression of Nlk significantly upregulated Pgrn levels in the medium in a kinase activity-dependent manner in BV2 microglial cells. Representative western blot images **(D)** and quantification **(E)** of Pgrn levels in BV2 cells by Nlk overexpression. Nlk-T298A is a kinase-inactive form of *Nlk.* *P < 0.05, **P < 0.01, ***P < 0.001, N.S., non-significant (*one-way ANOVA* with Tukey’s post-hoc testing, n=3 per group).

To determine if overexpression of Nlk, and the kinase activity of Nlk, affect the regulation of Pgrn expression, we transiently transfected Flag-tagged *Nlk* and examined the expression levels of Pgrn protein in BV2 cells (Figure 6D,E). Consistent with the loss-of-function studies, increased expression of wild-type Nlk (Nlk-WT) strongly increased extracellular, but not intracellular, Pgrn expression levels in BV2 cells (Figure 6D,E). In contrast, kinase-inactive Nlk (Nlk-T298A) had no effect on the regulation of Pgrn in BV2 cells. T298A is a threonine to alanine substitution at residue 298 in the catalytic domain of Nlk, leading to defective kinase activity. Taken together, these data strongly suggest that Nlk deficiency decreases, and its overexpression increases, Pgrn levels in a kinase activity-dependent manner in microglia both *in vivo* and *in vitro.*

### Nlk controls Pgrn endocytosis in microglia in a clathrin-dependent manner

We next sought to elucidate the cellular mechanism through which Nlk regulates Pgrn expression levels in microglia. We first examined the specific subcellular localization of Pgrn in WT and *Nlk^+/−^* primary microglia by immunofluorescence staining. We found that the expression of Pgrn in WT primary microglia was weak and mainly diffuse (Figure 7A and Figure 7-figure supplement 1). In contrast, the expression pattern of Pgrn in *Nlk^+/−^* primary microglia was dramatically punctate in morphology, leading to increased Pgrn fluorescence intensity in *Nlk^+/−^*microglia due to the more restricted, localized expression pattern (Figure 7A,B and Figure 7-figure supplement 1). This suggests that Pgrn may be trafficked into the endosome/lysosome-like vesicles in *Nlk^+/−^* primary microglia, since Pgrn is known to co-localize with lysosome-associated membrane protein 1 (Lamp1) in neurons (Hu et al., 2010; Tanaka et al., 2017). To test this idea, we co-stained for Pgrn and several lysosome and endosome markers in WT and *Nlk^+/−^* primary microglia. As expected, Pgrn partially co-localized with Lamp1- and Lamp2-positive lysosomes, as well as with early (Rab5), late (Rab7), and recycling (Rab11) endosomes, which were significantly increased in *Nlk^+/−^* primary microglia compared to WT littermate controls (Figure 7A and Figure 7-figure supplement 1), suggesting Nlk mediates trafficking of Pgrn into the endosome/lysosome pathway. Due to the lack of available antibodies for immunofluorescence detection of Nlk, we have not been able to demonstrate the subcellular co-compartmentalization of endogenous Nlk protein to similar intracellular structures as Pgrn.

**Figure 7.**
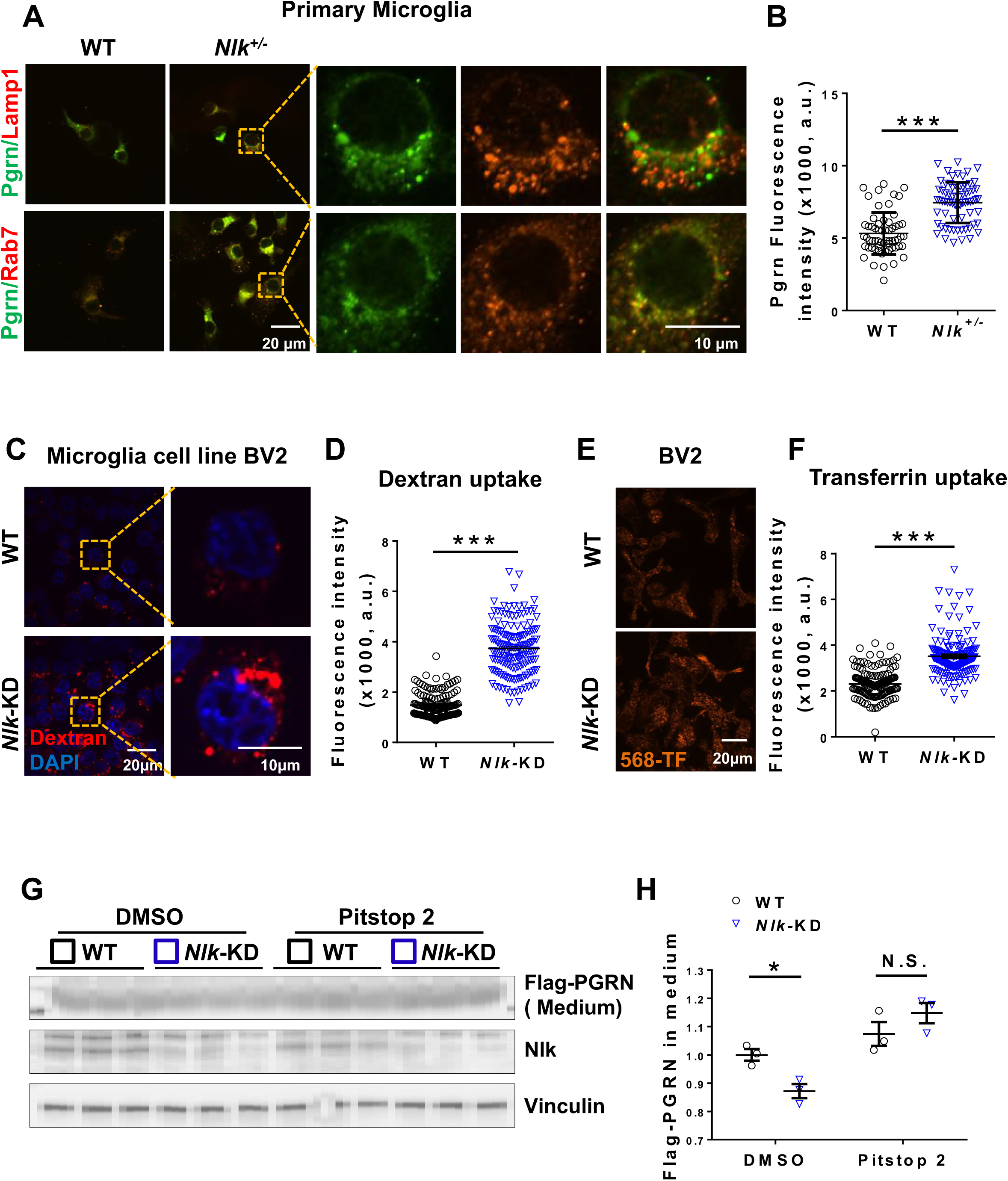
Nlk deficiency increases clathrin-dependent Pgrn endocytosis in microglia. **(A,B)** Nlk reduction resulted in the increased localization of Pgrn to lysosomes and late endosomes. **(A)** Representative images of WT and *Nlk^+/−^* primary microglia co-stained for Pgrn/Lamp1 (upper panels) or Pgrn/Rab7 (lower panels) demonstrated co-localization of Pgrn with lysosomes and late endosomes, respectively. Right panels are magnified views of areas marked in the left panels. **(B)** Quantification of WT and *Nlk^+/−^* primary microglia stained for Pgrn. ***p < 0.001 (non-parametric Mann-Whitney *t-test,* n= ~70 cells per group from three individual wells). **(C,D)** Enhanced uptake of the extracellularly provided Dextran in *Nlk* -KD BV2 cells. Representative images **(C)** and quantification **(D)** of WT and Nlk-KD BV2 cells after 20 minutes incubation with 647-Dextran. ***p < 0.001 (non-parametric Mann-Whitney *t-test,* n = ~150 cells per group from three individual wells). **(E,F)** Enhanced uptake of the extracellularly provided Transferrin in Nlk-KD BV2 cells. Representative images **(E)** and quantification **(F)** of WT and Nlk-KD BV2 cells after 15 minutes incubation with Alexa568-Transferrin. ***p < 0.001 (non-parametric Mann-Whitney *t-test,* n = ~150 cells per group from three individual wells). **(G,H)** Representative western blot images **(G)** and quantification **(H)** showing the enhanced clathrin-dependent endocytosis of Flag-tagged PGRN provided exogenously in BV2 microglial cells. Cells were treated with DMSO or the clathrin-dependent endocytosis inhibitor Pitstop2 for 1 hour and incubated with the recombinant Flag-PGRN (10nM) for 15 minutes. *P < 0.05, N.S., non-significant (*two-way ANOVA,* n=3).

Collectively, the knowledge of the high degree of co-localization of Pgrn with endosome markers in *Nlk^+/−^* mice (Figure 7A and Figure 7-figure supplement 1), the enhanced Pgrn levels in extracellular medium following Nlk overexpression in BV2 cells (Figure 6D,E), and the fact that Pgrn is a secreted protein that can be endocytosed and delivered to lysosomes (Hu et al., 2010), raise the possibility that Nlk regulates microglial endocytosis. Thus, we decided to test whether Nlk functionally affects endocytosis in microglia. To do so, WT and *Nlk*-KD BV2 microglial cells were incubated with fluorescently-labeled Dextran. As expected, Nlk depletion significantly enhanced the degree of fluorescently-labeled Dextran uptake into BV2 cells (Figure 7C,D). We also found a similar effect using the fluorescently-labeled Transferrin in *Nlk*-KD BV2 microglial cells and primary microglia from *Nlk^+/−^* mice (Figure 7E,F and Figure 7-figure supplement 2A). To determine if Nlk regulates endocytosis in neurons as well, primary cortical neurons were cultured from WT, *Nlk^+/−^,* and *Nlk^−/−^* mice and Transferrin uptake was examined. In stark contrast to our results from BV2 microglial cells and primary microglia, *Nlk* deletion did not affect the uptake of fluorescently-labeled Transferrin in primary neurons (Figure 7-figure supplement 2B), indicating that Nlk reduction enhances endocytosis activity in microglia, but not in neurons.

Next, to directly test if Nlk affects microglial endocytosis of Pgrn specifically, we treated WT and *Nlk*-KD BV2 microglial cells with exogenous Flag-tagged recombinant PGRN proteins. We measured Flag-tagged PGRN proteins remaining in the medium after a 15 minute incubation and found a significantly decreased level of Flag-tagged PGRN remaining in media from the *Nlk*-KD BV2 cells compared to that from WT controls (Figure 7G,H). Since endocytosis can be divided into phagocytosis, pinocytosis, and receptor-mediated endocytosis, we decided to investigate which specific pathway is likely regulated by Nlk. Using Pitstop 2, a clathrin-mediated endocytosis inhibitor, we found that Nlk-regulated endocytosis of Flag-tagged PGRN was completely blocked. Although we cannot exclude the possibility that Nlk regulates the exocytosis of PGRN, these data suggest that Nlk plays a role in receptor-mediated endocytosis of PGRN (Figure 7G,H).

### Nlk controls Pgrn levels via lysosome-dependent degradation in microglia

Considering our evidence that Nlk regulates Pgrn uptake via endocytosis in microglia (Figure 7G,H), it is likely that Pgrn is then degraded via lysosome-dependent degradation. To reconcile the fact that Nlk reduction promotes microglial Pgrn endocytosis, resulting in a transient increase in vesicular Pgrn staining (Figure 7A,B), and that we observe an overall reduction in intracellular and extracellular Pgrn protein levels in our compound heterozygote animals (Figure 3A-C), we examined the involvement of Nlk-mediated lysosomal degradation of Pgrn. We treated primary microglia with the lysosome inhibitor Bafilomycin A1 (BafA1), which blocks the final lysosome-mediated degradation step by inhibiting vATPase-mediated acidification of lysosomes, and examined the expression patterns of Pgrn in WT and *Nlk^+/−^* primary microglia. The increased number and intensity of Pgrn puncta in *Nlk^+/−^* primary microglia relative to WT controls was completely abrogated by the BafA1 treatment, which further enhanced intensity in both groups (Figure 8A,B). Furthermore, biochemical analysis confirmed that Nlk deficiency-mediated degradation of both endogenous Pgrn (Figure 8C,D) and exogenously provided Flag-tagged PGRN (Figure 8E,F) was indeed dependent on lysosomal activity in BV2 microglial cells. Once again, Nlk did not affect the degradation of the exogenously provided recombinant Flag-PGRN in primary cortical neurons (Figure 8-figure supplement 1). Taken together, these studies strongly suggest that Nlk regulates Pgrn levels in the brain in a lysosome-dependent manner through microglia, but not through neurons.

**Figure 8.**
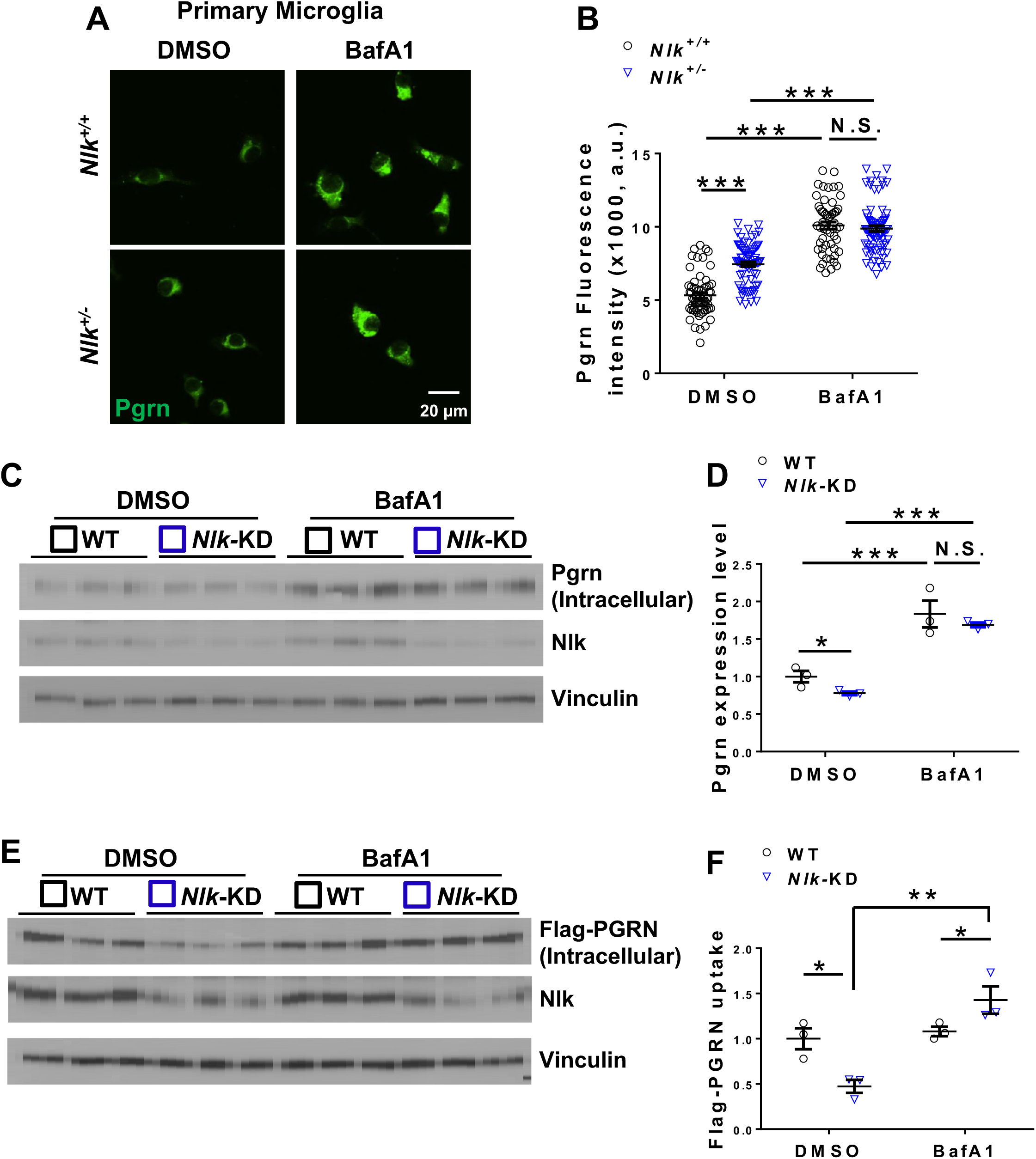
Nlk controls Pgrn degradation in microglia in a lysosome-dependent manner. **(A,B)** Representative images **(A)** and quantification **(B)** of WT and *Nlk^+/−^* primary microglia stained for Pgrn after 4 hours treatment with DMSO vesicle or the lysosomal inhibitor BafA1. ***p < 0.001, N.S., non-significant (*two-way ANOVA,* n = ~70 cells per group from three individual wells). **(C,D)** Representative western blot images **(C)** and quantification **(D)** showing the enhanced degradation of the endogenous Pgrn by Nlk reduction in BV2 microglial cells, which is dependent on lysosome activity. *p < 0.05, ***P < 0.001, N.S., non-significant (*two-way ANOVA,* n=3). **(E,F)** Representative western blot images **(E)** and quantification **(F)** showing the enhanced degradation of the exogenously provided recombinant Flag-PGRN protein in *Nlk-* KD BV2 cells, which is lysosome activity dependent. Cells were treated with DMSO or BafA1 for 4 hours and incubated with the recombinant Flag-PGRN (20nM) for 15 minutes. *P < 0.05, **P < 0.01 (*two-wayANOVA,* n=3).

## Discussion

Heterozygous and homozygous loss-of-function mutations in the *GRN* gene are common causes of familial FTLD and NCL, respectively (Baker et al., 2006; Cruts et al., 2006). Furthermore, reduced PGRN levels have been linked to other neurodegenerative disorders including AD (Jian et al., 2016; Perry et al., 2013; Wojtas et al., 2012). Despite extensive studies, the biological and molecular mechanisms underlying Pgrn function and metabolism have yet to be completely understood. Furthermore, there is no cure or effective therapeutic to reverse or slow progression of PGRN-associated neurodegenerative diseases. This underlines the need for basic research to better understand Pgrn neurobiology and to identify effective strategies or proteins that could increase Pgrn levels in the brain, which will eventually be useful for the development of effective therapeutics for several neurodegenerative disorders associated with PGRN levels. In this study, we identified such a protein, Nlk, that regulates Pgrn levels and the pathophysiological phenotypes of *Grn* haploinsufficiency. By combining mouse genetic approaches and cell biological studies, we show that Nlk is a key protein regulating Pgrn levels in the brain via control of the receptor-mediated endocytosis-lysosomal pathways in microglia, and thus modulates the pathophysiology of *Grn* haploinsufficiency in the brain (Figure 9).

**Figure 9.**
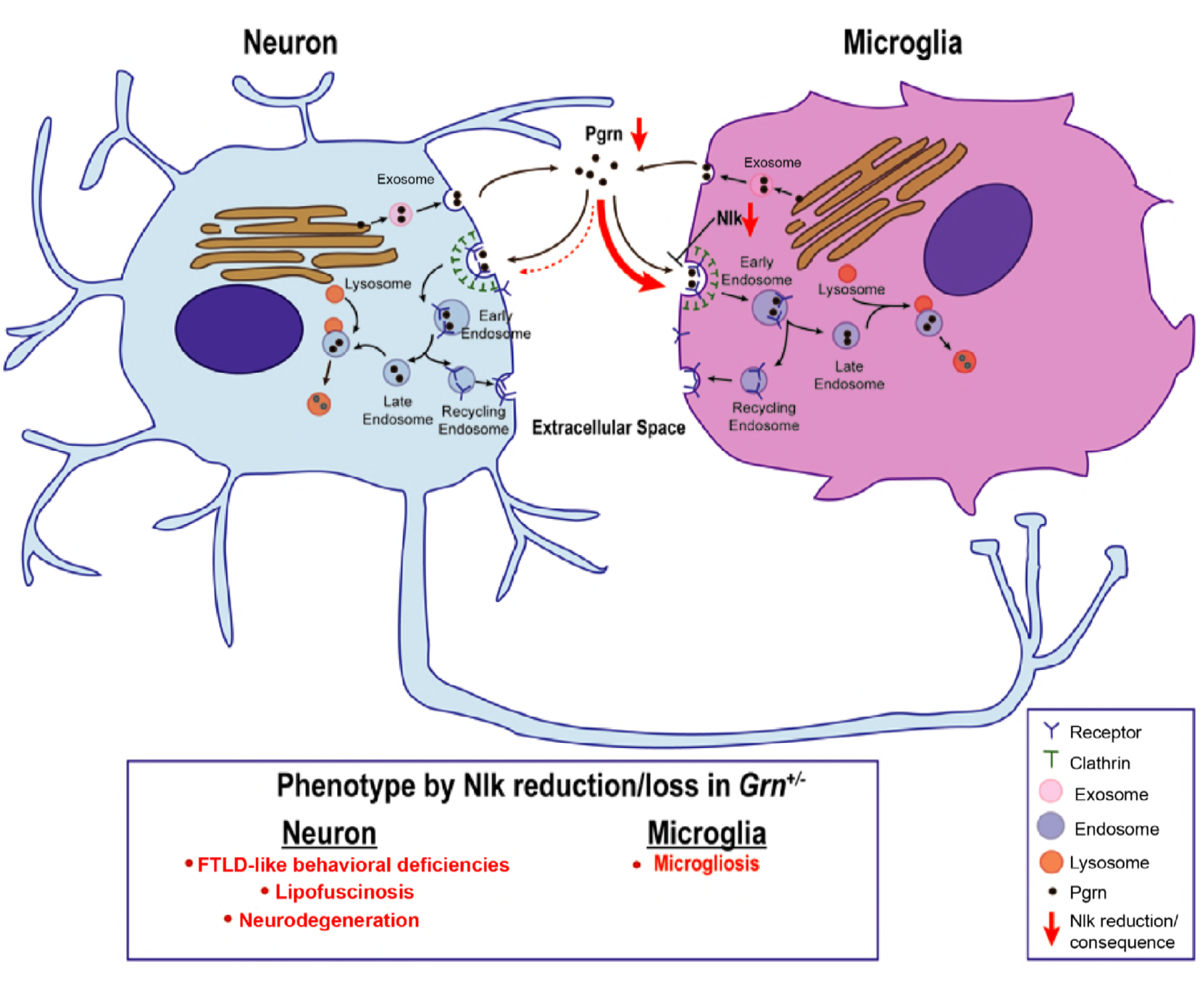
Model for the Nlk-mediated control of Pgrn levels through microglia, and the consequence of Nlk reduction in *Grn^+/−^* mice.

Different from *GRN* haploinsufficiency in humans, mice with a heterozygous loss of *Grn* fail to produce any obvious behavioral or neuropathological FTLD-PGRN phenotypes (Ahmed et al., 2010; Kayasuga et al., 2007). As a consequence, more attention has been directed towards characterizing mice that are null for *Grn,* which reproduces several features of human FTLD-PGRN and NCL (Filiano et al., 2013; Hafler et al., 2014; Klein et al., 2017; Yin et al., 2010b). Studies using *Grn^−/−^* mice would enhance our understanding of Pgrn function and biology in the brain, and provide some insights into the pathophysiology of FTLD-PGRN. However, there are clear limitations using *Grn* -deficient mice for such studies, one of which is whether and how increased expression levels of Pgrn would be able to rescue the phenotypes specific to FTLD-PGRN or NCL. Also, the lack of neuropathological phenotypes in *Grn^+/−^* heterozygous mice or the absence of *Grn* gene and Pgrn protein in *Grn^−/−^* null mice lead to an unequivocal desire for the development and identification of more physiologically relevant animal models for FTLD-PGRN study in order to better understand the molecular and cellular basis of neurodegeneration caused by *Grn* haploinsufficiency.

In this regard, it is very intriguing that this study shows that *Grn^+/−^* heterozygous mice develop several behavioral and neurodegenerative phenotypes, including cognitive impairments and altered anxiety, as well as lipofuscin accumulation, microgliosis, and retinal neurodegeneration, in the *Nlk^+/−^* heterozygous background (Figures 1,2), some of which are similarly observed in human patients with *GRN* haploinsufficiency (Viskontas et al., 2007; Ward et al., 2014; Woollacott and Rohrer, 2016). Consistent with previous studies, the *Grn^+/−^*heterozygous mice in the WT (*Nlk^+/+^*) background do not develop any phenotypes (Figures 1,2) (Ahmed et al., 2010; Kayasuga et al., 2007). To our knowledge, this is the first report showing that the genetically heterozygous *Grn^+/−^* mice can develop relevant cognitive impairments and neurodegeneration phenotypes in a certain genetic background, specifically a 50% Nlk expression reduction, which together leads to 70-80% Pgrn reduction (Figure 3A-C). Although several pathological phenotypes assessed in the present study do not quite reach statistical significance (p< 0.05) by *one-way ANOVA* in some brain regions, we believe that the statistically significant and trending data (significant by Mann-Whitney *t-test*) together represent a potential biologically significant threshold of required Pgrn levels below which pathological phenotypes begin to appear in mice. We speculate that the reduction of Pgrn by Nlk haploinsufficiency in the already *Grn* heterozygote lowers Pgrn protein levels to near threshold levels, and that perhaps a slight decrease in Pgrn dosage even further would generate significant pathological changes in all assessed brain regions.

*Grn* knockin mouse models have been generated with mutations found in human FTLD patients (PGRN-R504X and PGRN-R493X) (Fujita et al., 2018; Nguyen et al., 2018), and in particular, PGRN-R504X heterozygous mice exhibited Tau-mediated synaptic pathology and cognitive impairment, but not neurodegeneration (Fujita et al., 2018). Further studies are needed to investigate the phenotypic variance in mice of 50% Pgrn level in *Grn^+/−^* mice versus 50% Pgrn-R504X (with 50% WT Pgrn) level in Pgrn-R504X/+ heterozygous mice. Nevertheless, utilizing diverse *Grn* mutant mouse models and different genetic approaches will help us to better understand how *Grn* mutations affect the function and metabolism of Pgrn and its role in the pathogenesis of diverse neurodegenerative diseases including FTLD-PGRN.

The importance of expression levels and the functions of Pgrn in the brain, in particular in neurons and in microglia, have been well established (Kao et al., 2017; Petkau and Leavitt, 2014). A fundamental aim is to understand how Pgrn levels are regulated. In the brain, Pgrn protein is detected highly in neurons and activated microglia by immunostaining (Hu et al., 2010), while *Grn* mRNA is expressed highly in microglia and lowly in neurons and astrocytes (Zhang et al., 2014b). Conditional deletion of *Grn* specifically either in neurons or in microglia is not sufficient to cause phenotypes associated with Pgrn deficiency in mice (Petkau et al., 2017a; Petkau et al., 2017b), which may be due to extracellular Pgrn levels being maintained at functionally sufficient levels by different cell types. Our studies provide three significant findings on the regulation of Pgrn levels in the brain: (1) Nlk is critical for the regulation of Pgrn levels in the brain; (2) Nlk regulates Pgrn levels via microglia, but not via neurons; and (3) Nlk-mediated control of Pgrn levels occurs at the level of protein, but not the level of *Grn* mRNA expression. These findings support the main conclusion that Nlk controls Pgrn protein levels in the brain via microglia, which cannot be compensated for by other cell types. Consistent with this notion, Pgrn reduction levels (about 50% reduction) in the whole cortex of the microglia-specific KO (*Nlk^fl/fl^; Cx3cr1-cre*) mice (Figure 5A-C) are equivalent to those in Nlk deletion in all cell types (*Nlk^fl/fl^*; *Actin-cre^ERT2^*) (Figure 3F-H), suggesting that microglia are the major cell types by which Nlk plays a fundamental role in regulating Pgrn levels in the brain.

Another important outcome of this study is to reveal the cell biological mechanism underlying the Nlk-mediated regulation of Pgrn levels in microglia. Specifically, Nlk regulates Pgrn levels via control of the endocytosis-lysosomal pathways in microglia, but not in neurons: (1) Nlk negatively regulates microglial endocytosis (Figure 7); (2) Nlk negatively regulates receptor-mediated (or clathrin-dependent) PGRN endocytosis (Figure 7G,H); and (3) Nlk regulates PGRN levels via the lysosome-dependent degradation pathway (Figure 8). Together, this new information regarding the Nlk-mediated function in microglia on the regulation of Pgrn levels provides new cell biological regulatory mechanisms of Pgrn catabolism and its implication to the Pgrn biology and associated disease pathogenesis, highlighting the significance of the present study. It is unclear how Nlk regulates endocytosis and lysosome-dependent degradation. One possibility is that Nlk localizes into endosomes/lysosomes and regulates their dynamics. Although we have not successfully identified endogenous Nlk localization by antibody staining due to technical issues, it is worth noting that Nlk localizes to both the nucleus and cytoplasm when overexpressed in cell lines, without preferential localization to any specific organelle.

Several important questions are raised by our study. First, what is the molecular basis of the Nlk-mediated cell-type specific regulation of endocytosis? In other words, why does Nlk regulate endocytosis specifically in microglia, but not in neurons, and what are the direct substrate(s) of Nlk that control microglial endocytosis? Our preliminary studies suggest Nlk can phosphorylate several proteins involved in endocytosis, some of which are expressed specifically or very highly in microglia, but not in neurons. Second, what receptor mediates Pgrn endocytosis in microglia? Is it Sortlin since it is known to act as a Pgrn receptor for endocytosis and is able to regulate Pgrn levels (Hu et al., 2010)? Third, what effect does Nlk have on Pgrn secretion? Finally, considering the potential promising outcomes of Nlk overexpression in a microglia cell culture system (Figure 6D,E), can the increased expression of Nlk specifically in microglia upregulate Pgrn levels in the brain *in vivo* in order to rescue or improve the pathophysiology associated with reduced PGRN levels? Given that this regulation is kinase activity dependent, is Nlk a potential druggable target? Collectively, the findings of our study warrant a further, detailed mechanistic examination of the precise Nlk-mediated microglia-modulating effects, including characterization of the effects of varying Nlk levels on synaptic pruning and neuronal numbers in a brain circuit-specific manner, which may be involved in the behavioral changes we report here.

In conclusion, our study provides strong genetic and cell biological evidence that Nlk can control the microglial endocytosis-lysosomal pathway and modulate Pgrn levels and biology in the brain of a *Grn^+/−^* mouse model. It reveals a new regulatory mechanism for the control of Pgrn levels and, consequently, its associated neurobiology in the brain, which provide insight into the pathophysiology and the therapeutic development of PGRN-associated neurodegenerative diseases. This study also supports the future investigation of potential roles of Nlk and Nlk-mediated microglia modifications in diverse neurodegenerative diseases including AD.

## Methods

### Mouse husbandry and genetics

All animal experiments were approved by the Institutional Animal Care and Use Committee (IACUC) of Yale University and performed according to the National Institutes of Health guidelines for the Care and Use of Laboratory Animals. Mice were maintained on a 12 hour light/dark cycle with standard mouse chow and water *ad libitum.* Two distinct *Nlk* gene trap (RRJ297 and XN619) mouse lines were maintained on a pure C57BL/6J background (Ju et al., 2013). To generate *Nlk* heterozygous pups for primary microglia culture, *Nlk* heterozygous (*Nlk^RRJ297/+^* or *Nlk*^*XN619*/+^; simply *Nlk*^+/−^) mice and wild-type (WT) mice were crossed. To generate *Nlk* mutant pups for primary neuronal culture, *Nlk* heterozygous (RRJ297/+ or XN619/+) mice were intercrossed. All four F1 progenies, including WT (*Nlk*^+/+^), heterozygous (*Nlk^RRJ297/+^* or *Nlk^XN619/+^*; collectively *Nlk^+/−^*), and compound heterozygous (*Nlk^RRJ297/XN619^*; simply “knockout” (KO) here) pups, were used at postnatal day (P) 0. The *Nlk* conditional KO allele (*Nlk^fl/fl^*) were co-developed with Dr. Ernesto Canalis (Canalis et al., 2014) and deposited to the Jackson laboratory (JAX #024537). *Nlk^fl/fl^* mice were backcrossed onto a pure C57BL/6J background over 10 generations before mating to *Actin-cre^ERT2^* (Hayashi and McMahon, 2002), *Nex-cre* (Goebbels et al., 2006), or *Cx3cr1-cre* (Yona et al., 2013) mice. To activate the *cre* in *cre^ERT2^*, *Nlk^fl/fl^*; *Actin-cre^ERT2^* and their littermate control mice received daily intraperitoneal injections of tamoxifen (Sigma-Aldrich, T5648, 100mg/kg) for 7 consecutive days at 6 weeks of age. *Grn^−/−^* mice were obtained from the Jackson laboratory (JAX #013175). To perform the genetic interaction study, *Nlk^+/−^* heterozygote mice were bred to *Grn^+/−^* heterozygote mice and all four subsequent F1 progeny (WT, *Nlk^+/−^*, *Grn^+/−^*, and *Nlk^+/−^ Grn^+/−^*) were obtained. Both male and female mice were used in this study unless mentioned otherwise.

### Mouse behavioral tests

#### Elevated plus maze

Elevated plus maze was set at a height of 65cm and consisted of two open white Plexiglas arms, each arm 8cm wide x 30cm long and two enclosed arms (30cm x 5cm) with 15cm high walls, which were connected by a central platform (5cm x 5cm). Individual mice were placed at the center of the maze, facing one of the closed arms and observed for 5 minutes. All apparatus arms were cleaned with 70% ethanol after every trial. Data acquisition was recorded on a JVC Everio, G-series camcorder, and analysis of time spent on open and closed arms was performed using Panlab’s Smart tracking and analysis program, v2.5.

#### Morris Water Maze Test

Mice at 12 months were tested in a cylindrical tank of 100cm in diameter and 60cm in height. The tank was filled with water at 25°C and the platform was submerged 1cm below the water surface. The tank was divided into four quadrants with different navigational landmarks for each quadrant. The midpoint of the wall in each quadrant was used as the starting location from which animals were released into the water. In the hidden platform acquisition test, mice were allowed to swim freely to search for the escape platform within 60 seconds. The platform location remained constant throughout the test. The time taken to reach the platform was recorded as the escape latency. The mouse was kept on the platform for 10 seconds after it found the hidden platform. If mice failed to find the platform within 60 seconds, it was guided to the platform and placed on the platform for 10 seconds; in this case, the escape latency was recorded as 60 seconds for this trial. The same animal was released from a new starting quadrant 4 minutes after the previous trial. The experiment was repeated with 6 trials per mouse, with one trial every morning and afternoon for 3 consecutive days. The mean escape latency was measured to evaluate the spatial learning ability.

Twenty-four hours after the hidden platform acquisition test, probe trials were conducted by removing the platform. Mice were placed in the quadrant opposite to the removed hidden platform and were allowed to swim freely in the pool for 60 seconds. The percentage of time spent in the area around the original hidden platform was used to indicate long-term memory maintenance. At the end of the probe trial, a colorful flag was placed on the top of the hidden platform, which was opposite to the testing quadrant. The mouse was released from 3 different quadrants 7 times, and the time spent to find the flag was recorded as a measure of visual ability. The probe trials were recorded on a JVC Everio, G-series camcorder and analyzed using Panlab’s Smart tracking and analysis program, v2.5. The observer was blinded to genotype for the duration of behavioral testing.

### Western blot analysis

To examine protein expression levels, samples were prepared from the whole cortex of each mouse genotype or the cultured cells. To prepare extracellular protein samples from tissue, dissected mouse cortex was homogenized in PBS with a dounce homogenizer and centrifuged for 10 minutes at 6,000rpm at 4°C, with the resulting supernatant collected for extracellular proteins. Supernatant culture medium from BV2 and primary microglia was also collected for extracellular protein analysis as shown previously (Capell et al., 2011). To prepare intracellular protein samples, the pellet was lysed in lysis buffer (50mM Tris [pH7.5], 150mM NaCl, 0.1% SDS, 0.5% Triton X-100, 0.5% NP-40 and Roche complete protease inhibitor cocktail), rotated at 4°C for 20 minutes, and then centrifuged for 10 minutes at 13,000rpm at 4°C. The supernatant was collected as intracellular proteins. All samples were quantified and 20 or 40μg total protein from each sample was boiled for 10 minutes in sample buffer (BioRad #161-0737), loaded onto a SDS-PAGE, and transferred to a nitrocellulose membrane for western blot analysis. Membrane was blocked by incubation in 5% skimmed milk powder in TBST, incubated with primary antibodies in TBST containing 5% skimmed milk, then incubation in secondary antibodies conjugated with HRP, before detection with ECL reagents. The following primary antibodies were used: mouse anti-Vinculin (Sigma, V9264), sheep anti-Pgrn (R&D, AF2557), and rabbit anti-Nlk (Abcam, ab26050).

### Quantitative real-time reverse transcription polymerase chain reaction (qRT-PCR)

The RNA extraction, cDNA synthesis, and qRT-PCR were similarly performed as described (Kim et al., 2013). Total RNA was extracted from mouse cortex with the optional DNase digest step according to the manufacturer’s instructions (Qiagen, #74136). cDNA was synthesized using the iScript cDNA Synthesis Kit (Bio-Rad, #170-8891). qRT-PCR was performed using the C1000 Thermal Cycler and quantified using the CFX96 Real-Time System (Bio-Rad). The TaqMan gene expression assay and the iQ supermix (Bio-rad, #170-8862) were used for PCR amplification and real-time detection of PCR products. All RNA samples were analyzed in triplicate and normalized relative to *ACTB* expression levels. The following probes from Invitrogen were used: *Grn* (Mm00433848_m1), *Nlk* (Mm00476435_m1), and mouse *ACTB* (4352933E).

### Primary cortical neuron culture

Primary cortical neurons were prepared from neonatal pups at P0. Cerebral cortices were isolated and freed from meninges. Tissues were first digested with papain (Worthington, LS003126) and DNase I (Sigma, 10104159001) in HBSS (Gibco, 14170-112) for 30 minutes at 37°C, then washed three times with HBSS, and triturated with fire-polished glass pipettes until single cells were obtained. Cell suspensions were then filtered through a 40μm cell strainer and seeded at different densities according to the experimental design. Neurons were first plated in Neurobasal medium (Gibco, 21103-049) supplemented with B27 (Gibco, 17504-044) and 1% FBS (Gibco, 16140-071) and changed to non-serum medium 1 day later. Half media changes were performed every 3 days. Neurons were treated and collected at 7 to 8 days *in vitro* (DIV-7/8) for endocytosis analysis or collected at DIV-14 for protein and mRNA expression.

### Primary microglial cell culture

Primary cultured microglia were prepared from mouse brains, mainly from the cortex and the hippocampus, at P2 or P3. Meninges were removed mechanically, and the cells were dissociated by 0.25% Trypsin (Gibco, 25200-056) in HBSS, then cultured in PDL (Sigma, P6407)-coated T25 flask with DMEM (Gibco, 11965-092) supplemented with 10% heat-inactivated FBS (65°C, 30 minutes) and penicillin/streptomycin. After 14 days (DIV-14), the culture flasks were shaken at 200rpm for 3 hours to collect microglia. Microglia from three flasks having the same genotype were put together and counted as one sample. In total, three independent experiments were repeated. All experiments were performed at DIV-16 to DIV-18.

### Generation of CRISPR-mediated *Nlk* knockdown cells

The CRISPR/Cas9 technology (Cong et al., 2013) was used to generate *Nlk* knockdown (*Nlk-* KD) cells. Two guide RNAs were cloned separately into px462_v2 plasmid (Addgene, #62987, a gift from Dr. Feng Zhang). 250ng of each plasmid was co-transfected into the murine microglial cell line BV2 using Nucleofector Kits (Lonza, VPI-1006). At 24 hours post transfection, cells were enriched by the treatment of 4μg/ml puromycin (Thermo Fisher Scientific, #A1113802) for an additional 48 hours. Living cells were dissociated and diluted to 1 cell/100μl in DMEM and cultured in 96-well plate to obtain clonal population. Gene-targeting efficiency for each clone was analyzed by western blot analysis. Due to the incomplete targeting of *Nlk* genes on the non-diploid BV2 cancer cell line, a single *Nlk* KD BV2 cell clone was identified and selected for our analyses. The two guide RNA sequences used were, #1; 5’-CCCATCCCCGGCACCGGGTC - 3’ and #2; 5’-AACAACGGGTCCCAAATTGT −3’.

### Cell culture

The murine microglial BV2 cells were grown in DMEM supplemented with 10% heat-inactivated FBS and maintained at 37°C and 5% CO_2_. Cells were transiently transfected, using Amaxa Nucleofector^®^ (Lonza, Walkersville, MD) according to the manufacturer’s instructions, with the *pcDNA3.1, Flag-mNlk-WT* or *Flag-mNlk-T298A* constructs. For lysosomal-mediated degradation inhibition experiments, BV2 cells were grown in a 12-well plate to 70 to 80% confluence before treatment. Lysosomal degradation was blocked by Bafilomycin-A1 (LC laboratories, B-1080, 400nM) for 4 hours. To investigate the endocytosis-lysosome-mediated degradation of PGRN in WT or *Nlk*-KD BV2 cells, cells were treated first with BafA1 or DMSO for 4 hours, the culture medium was removed, and cells were washed twice with PBS before treating with 20nM of recombinant human PGRN (Flag-PGRN recombinant, AdipoGen, AG-40A-0068) for 15 minutes. For primary cortical neurons, the recombinant human PGRN incubation time was increased to 1 hour with a 20nM concentration. For the clathrin-mediated endocytosis study, cells were treated first with Pitstop2 (Sigma, SML1169, 30μΜ) or DMSO for 1 hour, then the culture medium was removed, and cells were treated with 10nM of recombinant human PGRN for 15 minutes before collecting medium.

### Dextran uptake

BV2 microglial cells were plated on 24-well plates with a round glass coverslip (Electron Microscopy Sciences, #72196-12) for 2 days prior to experiment. The cells were placed on ice for 10 minutes, followed by incubation with Alexa 647-conjugated Dextran (Molecular Probes, #D22914) at 1mg/ml in DMEM for 20 minutes at 37°C. After washing twice with PBS, cells were fixed with 4% paraformaldehyde (PFA, Sigma, 158127) for 10 minutes.

### Transferrin uptake

BV2 cells, primary microglia, or primary cortical neurons were plated on PDL-coated glass bottom dishes (MatTek, P35G-1.5-14-C) 2 days before experiments. Cells were placed on ice for 10 minutes, followed by incubation with Alexa 568-conjugated Transferrin (Molecular Probes, T23365) at 25μg/ml in Live Cell Imaging Solution (LCIS, Invitrogen, A14291DJ) containing 20mM Glucose and 1% BSA for 15 minutes at 37°C. Cells were washed twice with cold LCIS before live cell image acquisition.

### Immunofluorescence staining

Immunofluorescence staining and confocal microscopy analyses were performed using frozen mouse tissues or cultured cells as described previously (Ju et al., 2013; Todd et al., 2015). Briefly, mouse brains were carefully removed after cardiac perfusion, fixed overnight in 4% PFA, and incubated in 20% and 30% sucrose gradient per day at 4°C. Cultured cells were also fixed in 4% PFA for 10 minutes. Sectioned mouse brain slides or cells were permeablized in PBS with 0.5% Triton X-100, and incubated with a blocking buffer (5% normal goat serum and 0.05% Triton X-100 in PBS), followed by primary antibody incubation in the blocking buffer at 4°C overnight, and secondary antibody incubation for 2 hours. Brain sections were incubated with TO-PRO-3 Iodide (642/661) (Invitrogen, T3605) and mounted in Vectashield (Vector Laboratories, H1400). Cultured cells were directly mounted in Vectashield with DAPI (Vector Laboratories, H1500). The following primary antibodies were used in this study: sheep anti-Pgrn (R&D, AF2557), rabbit anti-Iba1 (Wako, #019-19741), rabbit anti-Rab5 (Cell Signaling, #3547), rabbit anti-Rab7 (Cell Signaling, #2094), rabbit anti-Rab11 (Cell Signaling, #5589), rat anti-CD68 (Abcam, ab53444), rat anti-Lamp1 (Developmental Studies Hybridoma Bank, 1D4B), and rat anti-Lamp2 (Developmental Studies Hybridoma Bank, ABL-93). Pgrn expression was detected in mouse brain sections with antigen retrieval (10mM Citrate pH6.0 for 20 minutes at 90°C) before primary antibody was applied.

### Autofluorescence analysis

For autofluorescence analysis on mouse brains, sectioned tissues were washed in PBS. Without primary or secondary antibody incubation, sectioned brain tissues were coverslipped with Vectashield with DAPI and visualized on a Zeiss LSM710 Spectral confocal microscope with multiple excitation wavelengths including 488 nm and 543 nm. Images are z-stack composites encompassing the entire section.

### Retinal ganglion cell counts

Retinas from both eyes were dissected and processed as described (Klein et al., 2017). Briefly, retinas were post-fixed for 1 to 3 hours in 4% PFA, cryoprotected in 30% sucrose solution overnight at 4°C, embedded using OCT, and rapidly frozen on dry ice. 15μm sections were sliced on a cryostat and collected immediately on microscope slides. Retinal sections were then stained on slides. Primary antibody (mouse anti-Brn3a, Santa Cruz, 14A6, sc-8429) was applied overnight at 4°C. The secondary antibody used was Alexa Fluor 488-conjugated goat anti-mouse IgG (Invitrogen, A11001). TO-PRO-3 Iodide (642/661) was used to stain nuclei. Fluorescent images were scanned using a Zeiss LSM710 Spectral confocal microscope with a 40X objective and processed with Zen 2.1 software (Carl Zeiss). Within the ganglion cell layer, the number of immunostained Brn3a cells was counted per 100mm for each retinal section from the central regions. Data were averaged from 3 slices of each retina.

### Image acquisition and data analyses

For cultured cells, images were captured using Zeiss spinning disk confocal microscopy (SDC) with a 60X oil objective lens. For mouse brain sections, images were captured using a Zeiss LSM710 Spectral confocal microscope with a 20X or 40X objective and processed with Zen 2.1 software (Carl Zeiss), followed by z-stack composites encompassing the entire section. Three sections per animal were analyzed. Quantification of fluorescence intensity was performed by Volocity software (PerkinElmer).

### Statistical analysis

Data are presented as mean ± standard errors of measurement (SEM). The statistical significance was assessed using the *t-test, one-way ANOVA,* or *two-way ANOVA* on the GraphPad Prism 6.0 software (GraphPad Software). A value of P< 0.05 was considered statistically significant.

### Data availability

The datasets generated during and/or analyzed in the current study are available in the main and supplementary figures, as well as available from the corresponding author on reasonable request.

## Acknowledgements

We would like to thank Dr. Klaus-Armin Nave for providing *Nex-cre* mice, Dr. Katerina Akassoglou for BV2 cells, Levi M. Smith for technical advice for mouse behavioral assays; Dr. Stephen M. Strittmatter, Dr. Pietro De Camilli and the members of the Lim laboratory for their thoughtful comments. This work was supported by the National Institute of Neurological Disorders and Stroke grants R01 NS083706 and R01 NS088321.

## Competing Interests

The authors have no competing interests to disclose at this time.

## Author Contributions

T.D. and J.L. conceived the project. T.D. and H.K. performed experiments. T.D., H.K., T.M.D., L.T. and J.L. analyzed and interpreted data, and wrote the manuscript. All authors reviewed the manuscript and discussed the work.

---

**Figure 1-figure supplement 1.**
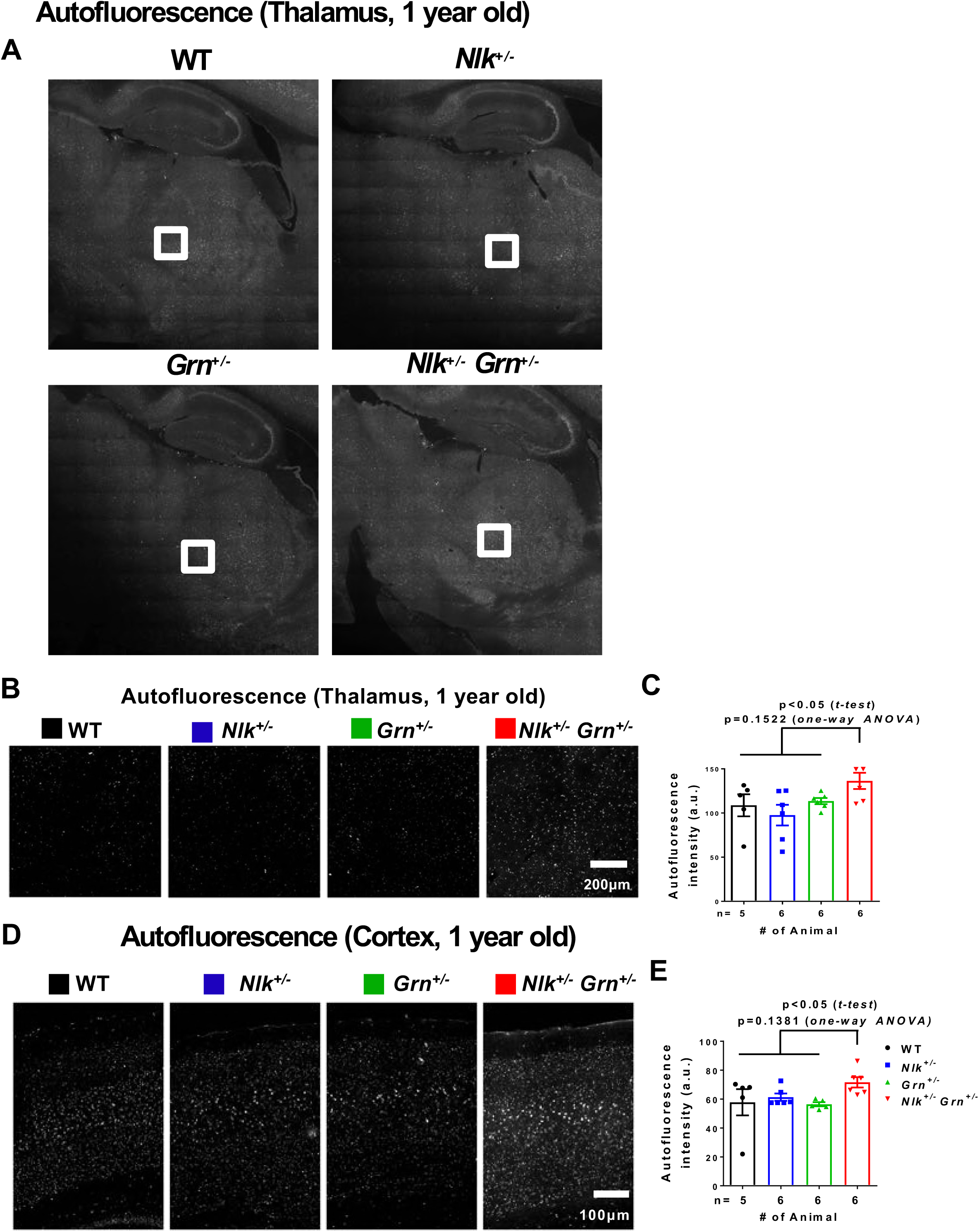
Enhanced accumulation of lipofuscin in the thalamus and cortex of *Nlk^+/−^ Grn^+/−^* mice. **(A-C)**Representative images **(A,B)** and quantification **(C)** of autofluorescence using 488 nm excitation in the thalamus of 1-year-old WT, *Nlk^+/−^*, *Grn^+/−^*, *Nlk^+/−^Grn^+/−^* mice. Areas marked by squares are magnified and seen in **(B).** P=0.1522 (*one-way NOVA* with Tukey’s multiple comparisons post-hoc test); *P < 0.05 (non-parametric Mann-Whitney *t-test).* **(D,E)** Representative images **(D)** and quantification **(E)** of autofluorescence in the cortex of 1-year-old WT, *Nlk^+/−^*, *Grn^+/−^*, *Nlk^+/−^Grn^+/−^* mice. P=0.1381 (*one-way ANOVA* with Tukey’s multiple comparisons post-hoc test); *P < 0.05 (non-parametric Mann-Whitney *t-test).*

**Figure 7-figure supplement 1.**
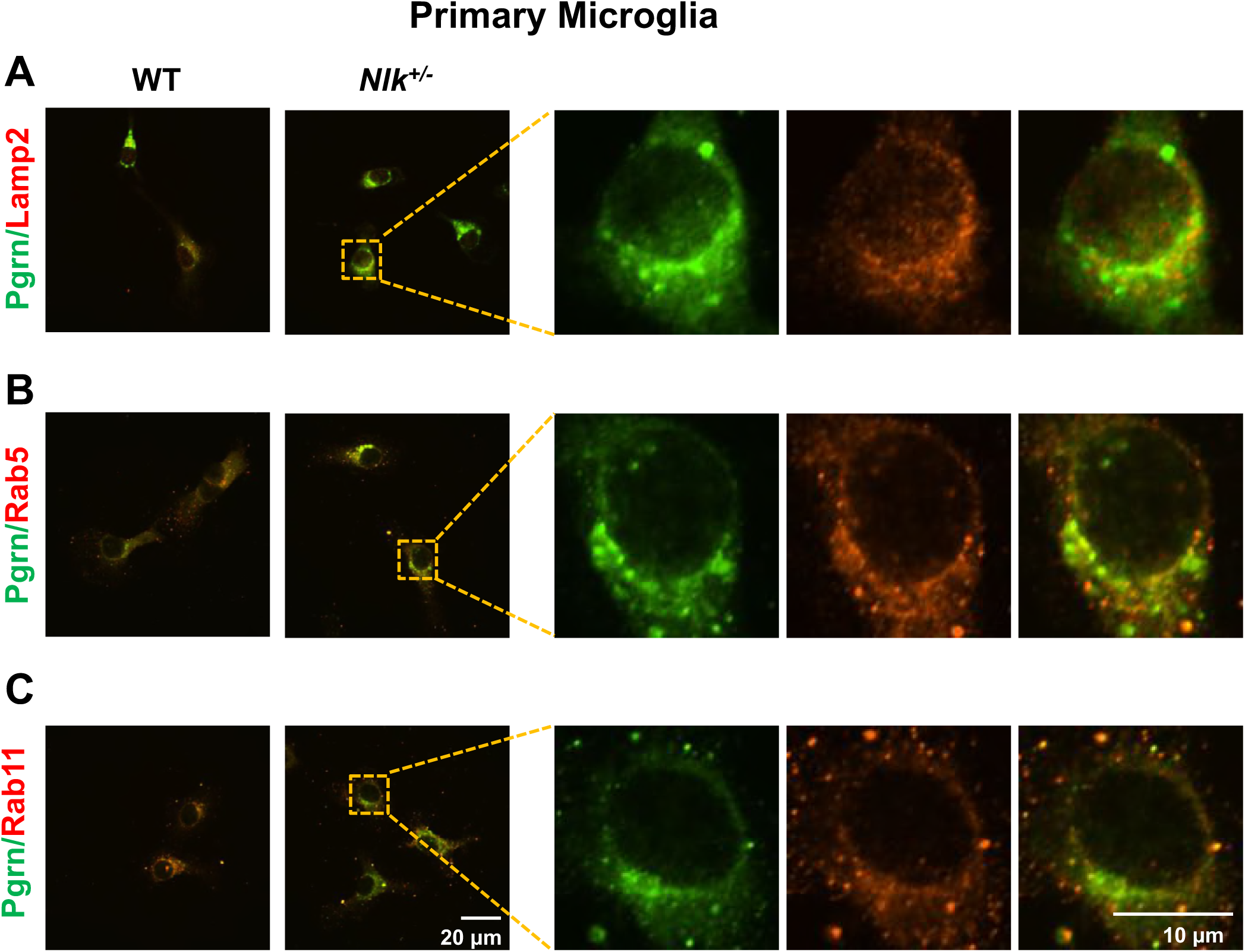
Nlk reduction leads to the increased localization of Pgrn to lysosomes and endosomes. Representative images of WT and *Nlk^+/−^* primary microglia co-stained for Pgrn/Lamp2 **(A),** Pgrn/Rab5 **(B),** and Pgrn/Rab11 **(C)** showing co-localization of Pgrn with lysosomes and early or recycling endosomes. Right panels are magnified views of areas marked in the left panels.

**Figure 7-figure supplement 2.**
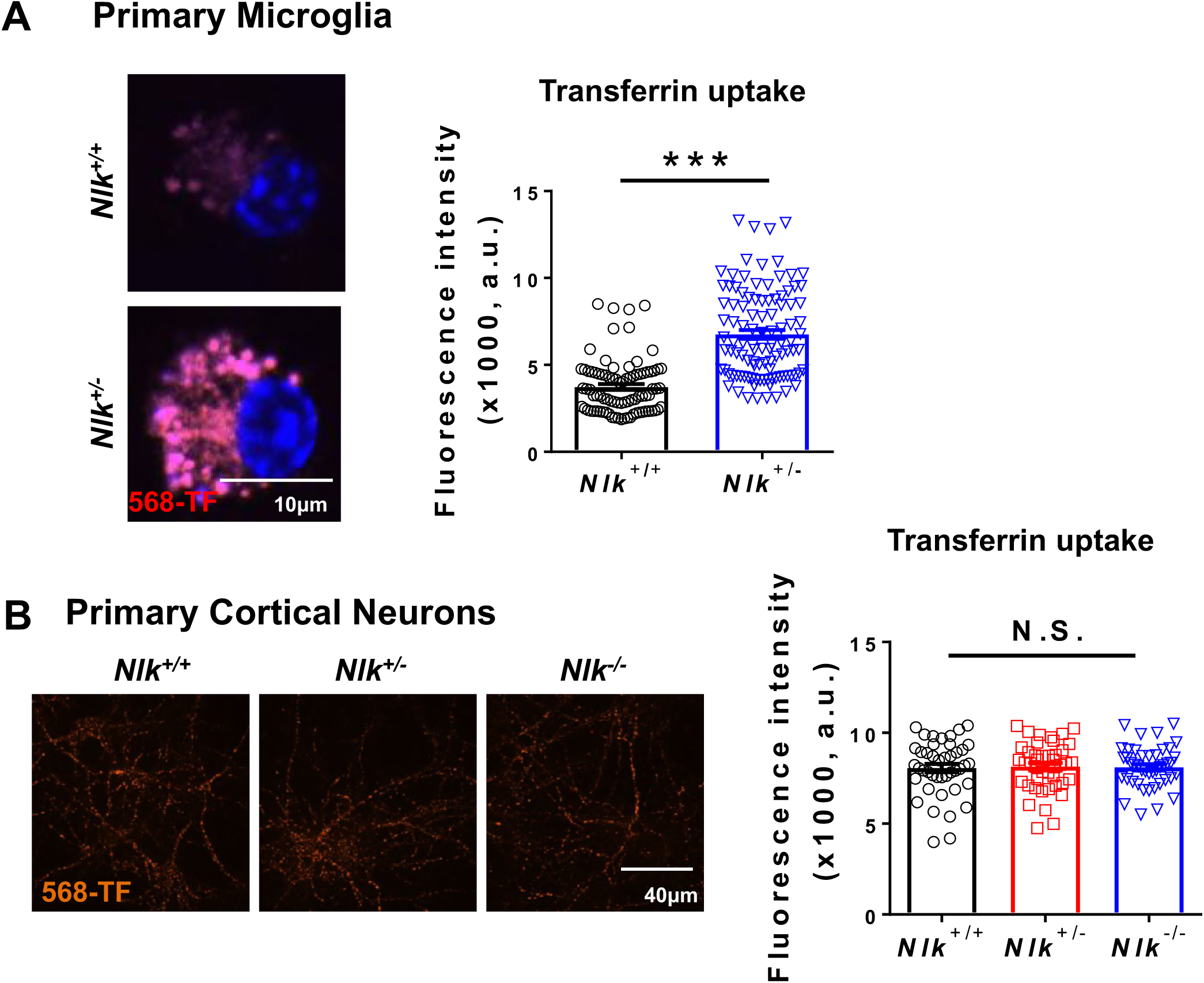
Nlk deficiency enhances endocytosis in microglia, but not in neurons. **(A)** Enhanced uptake of the extracellularly provided Transferrin in *Nlk^+/−^* primary microglia. Representative images (left panel) and quantification (right panel) of WT and *Nlk^+/−^*primary microglia after 15 minutes incubation with Alexa568-Transferrin. ***P < 0.001 (non-parametric Mann-Whitney *t-test,* n = ~100 cells measured per group from four individual wells). **(B)** No effects of Nlk on endocytosis in primary cortical neurons. Representative images (left panel) and quantification (right panel) of WT, *Nlk^+/−^*, *Nlk^−/−^* cortical neurons after 15 minutes incubation with Alexa568-Transferrin. N.S., non-significant (*one-way ANOVA* with Tukey’s multiple comparisons post-hoc test, n = ~50 cells measured per group from three individual wells).

**Figure 8-figure supplement 1.**
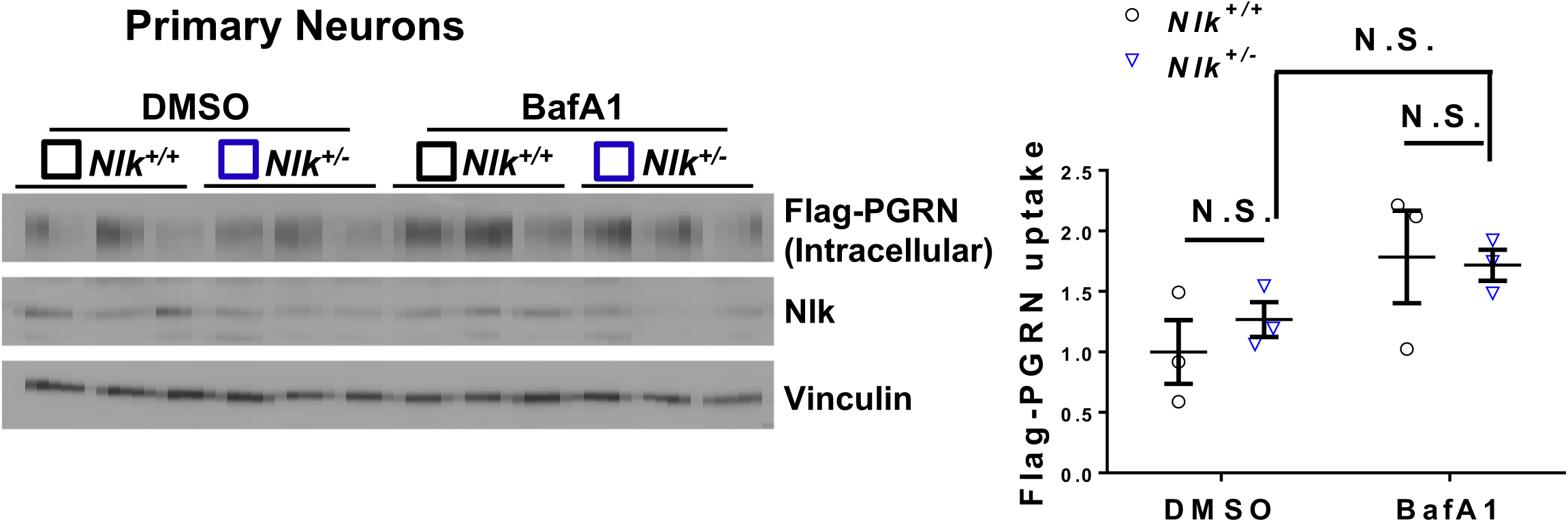
Nlk does not affect the expression levels of Pgrn provided extracellularly in primary cortical neurons. Representative western blot images (left panel) and quantification (right panel) showing that Nlk levels do not alter the lysosome-dependent degradation of exogenously provided recombinant Flag-PGRN protein in primary cortical neurons. WT and *Nlk^+/−^* primary cortical neurons were treated with DMSO or BafA1 for 4 hours and incubated with the recombinant Flag-PGRN (20nM) for 1 hour. N.S., non-significant (*two-way ANOVA*, n=3).

## References

1. Ahmed, Z., Mackenzie, I.R., Hutton, M.L., and Dickson, D.W. (2007). Progranulin in frontotemporal lobar degeneration and neuroinflammation. J Neuroinflammation 4, 7.

2. Ahmed, Z., Sheng, H., Xu, Y.F., Lin, W.L., Innes, A.E., Gass, J., Yu, X., Wuertzer, C.A., Hou, H., Chiba, S., et al. (2010). Accelerated lipofuscinosis and ubiquitination in granulin knockout mice suggest a role for progranulin in successful aging. Am J Pathol 177, 311–324.

3. Altmann, C., Vasic, V., Hardt, S., Heidler, J., Haussler, A., Wittig, I., Schmidt, M.H., and Tegeder, I. (2016). Progranulin promotes peripheral nerve regeneration and reinnervation: role of notch signaling. Mol Neurodegener 11, 69.

4. Baker, M., Mackenzie, I.R., Pickering-Brown, S.M., Gass, J., Rademakers, R., Lindholm, C., Snowden, J., Adamson, J., Sadovnick, A.D., Rollinson, S., et al. (2006). Mutations in progranulin cause tau-negative frontotemporal dementia linked to chromosome 17. Nature 442, 916–919.

5. Canalis, E., Kranz, L., and Zanotti, S. (2014). Nemo-like kinase regulates postnatal skeletal homeostasis. Journal of cellular physiology 229, 1736–1743.

6. Capell, A., Liebscher, S., Fellerer, K., Brouwers, N., Willem, M., Lammich, S., Gijselinck, I., Bittner, T., Carlson, A.M., Sasse, F., et al. (2011). Rescue of progranulin deficiency associated with frontotemporal lobar degeneration by alkalizing reagents and inhibition of vacuolar ATPase. J Neurosci 31, 1885–1894.

7. Cenik, B., Sephton, C.F., Kutluk Cenik, B., Herz, J., and Yu, G. (2012). Progranulin: a proteolytically processed protein at the crossroads of inflammation and neurodegeneration. J Biol Chem 287, 32298–32306.

8. Chen-Plotkin, A.S., Xiao, J., Geser, F., Martinez-Lage, M., Grossman, M., Unger, T., Wood, E.M., Van Deerlin, V.M., Trojanowski, J.Q., and Lee, V.M. (2010). Brain progranulin expression in GRN-associated frontotemporal lobar degeneration. Acta Neuropathol 119, 111–122.

9. Christopher, M.A., Myrick, D.A., Barwick, B.G., Engstrom, A.K., Porter-Stransky, K.A., Boss, J.M., Weinshenker, D., Levey, A.I., and Katz, D.J. (2017). LSD1 protects against hippocampal and cortical neurodegeneration. Nat Commun 8, 805.

10. Cong, L., Ran, F.A., Cox, D., Lin, S., Barretto, R., Habib, N., Hsu, P.D., Wu, X., Jiang, W., Marraffini, L.A., et al. (2013). Multiplex genome engineering using CRISPR/Cas systems. Science 339, 819–823.

11. Cruts, M., Gijselinck, I., van der Zee, J., Engelborghs, S., Wils, H., Pirici, D., Rademakers, R., Vandenberghe, R., Dermaut, B., Martin, J.J., et al. (2006). Null mutations in progranulin cause ubiquitin-positive frontotemporal dementia linked to chromosome 17q21. Nature 442, 920–924.

12. de Vugt, M.E., Riedijk, S.R., Aalten, P., Tibben, A., van Swieten, J.C., and Verhey, F.R. (2006). Impact of behavioural problems on spousal caregivers: a comparison between Alzheimer’s disease and frontotemporal dementia. Dement Geriatr Cogn Disord 22, 35–41.

13. Filiano, A.J., Martens, L.H., Young, A.H., Warmus, B.A., Zhou, P., Diaz-Ramirez, G., Jiao, J., Zhang, Z., Huang, E.J., Gao, F.B., et al. (2013). Dissociation of frontotemporal dementia-related deficits and neuroinflammation in progranulin haploinsufficient mice. J Neurosci 33, 5352–5361.

14. Fujita, K., Chen, X., Homma, H., Tagawa, K., Amano, M., Saito, A., Imoto, S., Akatsu, H., Hashizume, Y., Kaibuchi, K., et al. (2018). Targeting Tyro3 ameliorates a model of PGRN-mutant FTLD-TDP via tau-mediated synaptic pathology. Nat Commun 9, 433.

15. Gass, J., Lee, W.C., Cook, C., Finch, N., Stetler, C., Jansen-West, K., Lewis, J., Link, C.D., Rademakers, R., Nykjaer, A., et al. (2012). Progranulin regulates neuronal outgrowth independent of sortilin. Mol Neurodegener 7, 33.

16. Ghoshal, N., Dearborn, J.T., Wozniak, D.F., and Cairns, N.J. (2012). Core features of frontotemporal dementia recapitulated in progranulin knockout mice. Neurobiol Dis 45, 395–408.

17. Gijselinck, I., Van Broeckhoven, C., and Cruts, M. (2008). Granulin mutations associated with frontotemporal lobar degeneration and related disorders: an update. Hum Mutat 29, 1373–1386.

18. Goebbels, S., Bormuth, I., Bode, U., Hermanson, O., Schwab, M.H., and Nave, K.A. (2006). Genetic targeting of principal neurons in neocortex and hippocampus of NEX-Cre mice. Genesis 44, 611–621.

19. Hafler, B.P., Klein, Z.A., Jimmy Zhou, Z., and Strittmatter, S.M. (2014). Progressive retinal degeneration and accumulation of autofluorescent lipopigments in Progranulin deficient mice. Brain Res 1588, 168–174.

20. Hayashi, S., and McMahon, A.P. (2002). Efficient recombination in diverse tissues by a tamoxifen-inducible form of Cre: a tool for temporally regulated gene activation/inactivation in the mouse. Dev Biol 244, 305–318.

21. Hu, F., Padukkavidana, T., Vaegter, C.B., Brady, O.A., Zheng, Y., Mackenzie, I.R., Feldman, H.H., Nykjaer, A., and Strittmatter, S.M. (2010). Sortilin-mediated endocytosis determines levels of the frontotemporal dementia protein, progranulin. Neuron 68, 654–667.

22. Ishitani, T., Ishitani, S., Matsumoto, K., and Itoh, M. (2009). Nemo-like kinase is involved in NGF-induced neurite outgrowth via phosphorylating MAP1B and paxillin. J Neurochem 111, 1104–1118.

23. Ishitani, T., Ninomiya-Tsuji, J., Nagai, S., Nishita, M., Meneghini, M., Barker, N., Waterman, M., Bowerman, B., Clevers, H., Shibuya, H., et al. (1999). The TAK1 -NLK-MAPK-related pathway antagonizes signalling between beta-catenin and transcription factor TCF. Nature 399, 798–802.

24. Jian, J., Zhao, S., Tian, Q.Y., Liu, H., Zhao, Y., Chen, W.C., Grunig, G., Torres, P.A., Wang, B.C., Zeng, B., et al. (2016). Association Between Progranulin and Gaucher Disease. EBioMedicine 11, 127–137.

25. Ju, H., Kokubu, H., Todd, T.W., Kahle, J.J., Kim, S., Richman, R., Chirala, K., Orr, H.T., Zoghbi, H.Y., and Lim, J. (2013). Polyglutamine Disease Toxicity Is Regulated by Nemo-like Kinase in Spinocerebellar Ataxia Type 1. J Neurosci 33, 9328–9336.

26. Kao, A.W., McKay, A., Singh, P.P., Brunet, A., and Huang, E.J. (2017). Progranulin, lysosomal regulation and neurodegenerative disease. Nat Rev Neurosci 18, 325–333.

27. Kayasuga, Y., Chiba, S., Suzuki, M., Kikusui, T., Matsuwaki, T., Yamanouchi, K., Kotaki, H., Horai, R., Iwakura, Y., and Nishihara, M. (2007). Alteration of behavioural phenotype in mice by targeted disruption of the progranulin gene. Behav Brain Res 185, 110–118.

28. Kessenbrock, K., Frohlich, L., Sixt, M., Lammermann, T., Pfister, H., Bateman, A., Belaaouaj, A., Ring, J., Ollert, M., Fassler, R., et al. (2008). Proteinase 3 and neutrophil elastase enhance inflammation in mice by inactivating antiinflammatory progranulin. J Clin Invest 118, 2438–2447.

29. Kim, S., Chahrour, M., Ben-Shachar, S., and Lim, J. (2013). Ube3a/E6AP is involved in a subset of MeCP2 functions. Biochem Biophys Res Commun 437, 67–73.

30. Klein, Z.A., Takahashi, H., Ma, M., Stagi, M., Zhou, M., Lam, T.T., and Strittmatter, S.M. (2017). Loss of TMEM106B Ameliorates Lysosomal and Frontotemporal Dementia-Related Phenotypes in Progranulin-Deficient Mice. Neuron 95, 281–296.e6.

31. Lodato, S., and Arlotta, P. (2015). Generating neuronal diversity in the mammalian cerebral cortex. Annu Rev Cell Dev Biol 31, 699–720.

32. Lui, H., Zhang, J., Makinson, S.R., Cahill, M.K., Kelley, K.W., Huang, H.Y., Shang, Y., Oldham, M.C., Martens, L.H., Gao, F., et al. (2016). Progranulin Deficiency Promotes Circuit-Specific Synaptic Pruning by Microglia via Complement Activation. Cell 165, 921–935.

33. Meneghini, M.D., Ishitani, T., Carter, J.C., Hisamoto, N., Ninomiya-Tsuji, J., Thorpe, C.J., Hamill, D.R., Matsumoto, K., and Bowerman, B. (1999). MAP kinase and Wnt pathways converge to downregulate an HMG-domain repressor in Caenorhabditis elegans. Nature 399, 793–797.

34. Minami, S.S., Min, S.W., Krabbe, G., Wang, C., Zhou, Y., Asgarov, R., Li, Y., Martens, L.H., Elia, L.P., Ward, M.E., et al. (2014). Progranulin protects against amyloid beta deposition and toxicity in Alzheimer’s disease mouse models. Nat Med 20, 1157–1164.

35. Nguyen, A.D., Nguyen, T.A., Zhang, J., Devireddy, S., Zhou, P., Karydas, A.M., Xu, X., Miller, B.L., Rigo, F., Ferguson, S.M., et al. (2018). Murine knockin model for progranulin-deficient frontotemporal dementia with nonsense-mediated mRNA decay. Proc Natl Acad Sci U S A.

36. Ota, S., Ishitani, S., Shimizu, N., Matsumoto, K., Itoh, M., and Ishitani, T. (2012). NLK positively regulates Wnt/beta-catenin signalling by phosphorylating LEF1 in neural progenitor cells. Embo J 31, 1904–1915.

37. Perry, D.C., Lehmann, M., Yokoyama, J.S., Karydas, A., Lee, J.J., Coppola, G., Grinberg, L.T., Geschwind, D., Seeley, W.W., Miller, B.L., et al. (2013). Progranulin mutations as risk factors for Alzheimer disease. JAMA Neurol 70, 774–778.

38. Petkau, T.L., Blanco, J., and Leavitt, B.R. (2017a). Conditional loss of progranulin in neurons is not sufficient to cause neuronal ceroid lipofuscinosis-like neuropathology in mice. Neurobiol Dis 106, 14–22.

39. Petkau, T.L., Kosior, N., de Asis, K., Connolly, C., and Leavitt, B.R. (2017b). Selective depletion of microglial progranulin in mice is not sufficient to cause neuronal ceroid lipofuscinosis or neuroinflammation. J Neuroinflammation 14, 225.

40. Petkau, T.L., and Leavitt, B.R. (2014). Progranulin in neurodegenerative disease. Trends Neurosci 37, 388–398.

41. Petkau, T.L., Neal, S.J., Milnerwood, A., Mew, A., Hill, A.M., Orban, P., Gregg, J., Lu, G., Feldman, H.H., Mackenzie, I.R., et al. (2012). Synaptic dysfunction in progranulin-deficient mice. Neurobiol Dis 45, 711–722.

42. Petkau, T.L., Neal, S.J., Orban, P.C., MacDonald, J.L., Hill, A.M., Lu, G., Feldman, H.H., Mackenzie, I.R., and Leavitt, B.R. (2010). Progranulin expression in the developing and adult murine brain. J Comp Neurol 518, 3931–3947.

43. Raitano, S., Ordovas, L., De Muynck, L., Guo, W., Espuny-Camacho, I., Geraerts, M., Khurana, S., Vanuytsel, K., Toth, B.I., Voets, T., et al. (2015). Restoration of progranulin expression rescues cortical neuron generation in an induced pluripotent stem cell model of frontotemporal dementia. Stem Cell Reports 4, 16–24.

44. Rocheleau, C.E., Yasuda, J., Shin, T.H., Lin, R., Sawa, H., Okano, H., Priess, J.R., Davis, R.J., and Mello, C.C. (1999). WRM-1 activates the LIT-1 protein kinase to transduce anterior/posterior polarity signals in C. elegans. Cell 97, 717–726.

45. Rosen, E.Y., Wexler, E.M., Versano, R., Coppola, G., Gao, F., Winden, K.D., Oldham, M.C., Martens, L.H., Zhou, P., Farese, R.V., Jr., et al. (2011). Functional genomic analyses identify pathways dysregulated by progranulin deficiency, implicating Wnt signaling. Neuron 71, 1030–1042.

46. Rottinger, E., Croce, J., Lhomond, G., Besnardeau, L., Gache, C., and Lepage, T. (2006). Nemo-like kinase (NLK) acts downstream of Notch/Delta signalling to downregulate TCF during mesoderm induction in the sea urchin embryo. Development 133, 4341–4353.

47. Saijo, K., Winner, B., Carson, C.T., Collier, J.G., Boyer, L., Rosenfeld, M.G., Gage, F.H., and Glass, C.K. (2009). A Nurr1/CoREST pathway in microglia and astrocytes protects dopaminergic neurons from inflammation-induced death. Cell 137, 47–59.

48. Smith, K.R., Damiano, J., Franceschetti, S., Carpenter, S., Canafoglia, L., Morbin, M., Rossi, G., Pareyson, D., Mole, S.E., Staropoli, J.F., et al. (2012). Strikingly different clinicopathological phenotypes determined by progranulin-mutation dosage. Am J Hum Genet 90, 1102–1107.

49. Takahashi, H., Klein, Z.A., Bhagat, S.M., Kaufman, A.C., Kostylev, M.A., Ikezu, T., Strittmatter, S.M., and Alzheimer’s Disease Neuroimaging, I. (2017). Opposing effects of progranulin deficiency on amyloid and tau pathologies via microglial TYROBP network. Acta Neuropathol 133, 785–807.

50. Tanaka, Y., Suzuki, G., Matsuwaki, T., Hosokawa, M., Serrano, G., Beach, T.G., Yamanouchi, K., Hasegawa, M., and Nishihara, M. (2017). Progranulin regulates lysosomal function and biogenesis through acidification of lysosomes. Hum Mol Genet 26, 969–988.

51. Todd, T.W., Kokubu, H., Miranda, H.C., Cortes, C.J., La Spada, A.R., and Lim, J. (2015). Nemo-like kinase is a novel regulator of spinal and bulbar muscular atrophy. Elife 4, e08493.

52. Toh, H., Chitramuthu, B.P., Bennett, H.P., and Bateman, A. (2011). Structure, function, and mechanism of progranulin; the brain and beyond. J Mol Neurosci 45, 538–548.

53. Van Damme, P., Van Hoecke, A., Lambrechts, D., Vanacker, P., Bogaert, E., van Swieten, J., Carmeliet, P., Van Den Bosch, L., and Robberecht, W. (2008). Progranulin functions as a neurotrophic factor to regulate neurite outgrowth and enhance neuronal survival. J Cell Biol 181, 37–41.

54. Van Kampen, J.M., Baranowski, D., and Kay, D.G. (2014). Progranulin gene delivery protects dopaminergic neurons in a mouse model of Parkinson’s disease. PLoS One 9, e97032.

55. Viskontas, I.V., Possin, K.L., and Miller, B.L. (2007). Symptoms of frontotemporal dementia provide insights into orbitofrontal cortex function and social behavior. Ann N Y Acad Sci 1121, 528–545.

56. Ward, M.E., Taubes, A., Chen, R., Miller, B.L., Sephton, C.F., Gelfand, J.M., Minami, S., Boscardin, J., Martens, L.H., Seeley, W.W., et al. (2014). Early retinal neurodegeneration and impaired Ran-mediated nuclear import of TDP-43 in progranulin-deficient FTLD. J Exp Med 211, 1937–1945.

57. Wojtas, A., Heggeli, K.A., Finch, N., Baker, M., Dejesus-Hernandez, M., Younkin, S.G., Dickson, D.W., Graff-Radford, N.R., and Rademakers, R. (2012). C9ORF72 repeat expansions and other FTD gene mutations in a clinical AD patient series from Mayo Clinic. Am J Neurodegener Dis 1, 107–118.

58. Woollacott, I.O., and Rohrer, J.D. (2016). The clinical spectrum of sporadic and familial forms of frontotemporal dementia. J Neurochem 138 Suppl 1, 6–31.

59. Yin, F., Banerjee, R., Thomas, B., Zhou, P., Qian, L., Jia, T., Ma, X., Ma, Y., Iadecola, C., Beal, M.F., et al. (2010a). Exaggerated inflammation, impaired host defense, and neuropathology in progranulin-deficient mice. J Exp Med 207, 117–128.

60. Yin, F., Dumont, M., Banerjee, R., Ma, Y., Li, H., Lin, M.T., Beal, M.F., Nathan, C., Thomas, B., and Ding, A. (2010b). Behavioral deficits and progressive neuropathology in progranulin-deficient mice: a mouse model of frontotemporal dementia. FASEB J 24, 4639–4647.

61. Yona, S., Kim, K.W., Wolf, Y., Mildner, A., Varol, D., Breker, M., Strauss-Ayali, D., Viukov, S., Guilliams, M., Misharin, A., et al. (2013). Fate mapping reveals origins and dynamics of monocytes and tissue macrophages under homeostasis. Immunity 38, 79–91.

62. Zhang, H.H., Li, S.Z., Zhang, Z.Y., Hu, X.M., Hou, P.N., Gao, L., Du, R.L., and Zhang, X.D. (2014a). Nemo-like kinase is critical for p53 stabilization and function in response to DNA damage. Cell Death Differ 21, 1656–1663.

63. Zhang, Y., Chen, K., Sloan, S.A., Bennett, M.L., Scholze, A.R., O’Keeffe, S., Phatnani, H.P., Guarnieri, P., Caneda, C., Ruderisch, N., et al. (2014b). An RNA-sequencing transcriptome and splicing database of glia, neurons, and vascular cells of the cerebral cortex. J Neurosci 34, 11929–11947.

64. Zhou, X., Sun, L., Bastos de Oliveira, F., Qi, X., Brown, W.J., Smolka, M.B., Sun, Y., and Hu, F. (2015). Prosaposin facilitates sortilin-independent lysosomal trafficking of progranulin. J Cell Biol 210, 991–1002.

